# The *trans*-zeatin-type side-chain modification of cytokinins controls rice growth

**DOI:** 10.1101/2022.09.25.509405

**Authors:** Takatoshi Kiba, Kahori Mizutani, Aimi Nakahara, Yumiko Takebayashi, Mikiko Kojima, Tokunori Hobo, Yuriko Osakabe, Keishi Osakabe, Hitoshi Sakakibara

**Affiliations:** Graduate School of Bioagricultural Sciences, Nagoya University, Furocho, Chikusa-ku, Nagoya, 464-8601, Japan; RIKEN Center for Sustainable Resource Science, 1-7-22, Suehiro, Tsurumi, Yokohama 230-0045, Japan; Biosci. Biotech Center, Nagoya University., Furocho, Chikusa-ku, Nagoya, 464-8601, Japan; Department of Life Science and Technology, Tokyo Institute of Technology, Yokohama, 226-8501, Japan; Graduate School of Technology, Industrial and Social Sciences, Tokushima University, Tokushima, 770-8503, Japan

## Abstract

Cytokinins (CKs), a class of phytohormones with vital roles in growth and development, occur naturally with various side-chain structures, including *N*^*6*^-(Δ^2^-isopentenyl)adenine-, *cis*-zeatin- and *trans*-zeatin (tZ)-types. Recent studies in a model dicot plant *Arabidopsis* demonstrated that tZ-type CKs are biosynthesized via cytochrome P450 monooxygenase (P450) CYP735A, and have a specific function in shoot growth promotion. Although the function of some of these CKs has been demonstrated in a few dicotyledonous plant species, the significance of these variations and their biosynthetic mechanism and function in monocots and in plants with distinctive side-chain profiles than *Arabidopsis*, such as *Oryza sativa* (rice), remain elusive. In this study, we characterized *CYP735A3* and *CYP735A4* to investigate the role of tZ-type CKs in rice. Complementation test of the *Arabidopsis* CYP735A-deficient mutant and CK profiling of loss-of-function rice mutant, *cyp735a3 cyp735a4*, demonstrated that *CYP735A3* and *CYP735A4* encode P450s required for tZ-type side-chain modification in rice. *CYP735As* are expressed in both roots and shoots. The *cyp735a3 cyp735a4* mutants exhibited growth retardation concomitant with reduction in CK activity in both roots and shoots, indicating that tZ-type CKs function in growth promotion of both organs. Expression analysis revealed that tZ-type CK biosynthesis is negatively regulated by auxin, abscisic acid, and cytokinin and positively by dual nitrogen nutrient signals, namely glutamine-related and nitrate-specific signals. These results suggest that the physiological role of tZ-type CKs in rice is different from that in *Arabidopsis* and they control growth of both roots and shoots in response to internal and environmental cues in rice.

## Introduction

Cytokinins (CKs), a group of plant hormones, are involved in the regulation of various aspects of plant growth and development, including cell division and differentiation, organogenesis, root and shoot growth, and environmental responses (Mok and Mok, 2001; Sakakibara, 2006; Werner and Schmülling, 2009; Kiba et al., 2011; Kieber and Schaller, 2018; Kiba et al., 2019; Sakakibara, 2021). Natural CKs are mostly derivatives of adenine with a prenyl side chain at the *N*^6^ position. There are variations in the side-chain structure of CKs and the most common structures in plants are *N*^*6*^-(Δ^2^-isopentenyl)adenine (iP)-, *trans*-zeatin (tZ)-, and *cis*-zeatin (cZ)-types, which differ in the presence and stereoisomeric position of the terminal hydroxyl group on the prenyl side chain (Shaw, 1994; Mok and Mok, 2001; Gajdošová et al., 2011).

Our understanding of biosynthetic pathways of the side-chain variants has greatly progressed, thanks mostly to studies in *Arabidopsis*. The initial step of *de novo* iP- and tZ-type CK biosynthesis is catalyzed by adenosine phosphate-isopentenyltransferase (IPT) to produce iP-type CK nucleotide precursors, iP ribotides (iPRPs) (Kakimoto, 2001; Takei et al., 2001a). tZ-type side chain is formed by the enzyme cytochrome P450 monooxygenase CYP735A, which is encoded by *CYP735A1* and *CYP735A2* in *Arabidopsis* (Takei et al., 2004; Kiba et al., 2013). CYP735A *trans*-hydroxylates the prenyl side chain of iPRPs to produce tZ ribotides (tZRPs). Finally, the cytokinin-activating enzyme LONELY GUY (LOG) converts the nucleotide precursors to their active nucleobase forms, iP and tZ (Kurakawa et al., 2007; Kuroha et al., 2009; Tokunaga et al., 2012). The cZ-type CKs are biosynthesized via the prenylation of tRNA by tRNA-isopentenyltransferase (tRNA-IPT) and the degradation of the prenylated tRNA (Miyawaki et al., 2006; Gajdošová et al., 2011). Genes responsible for prenylated tRNA degradation for cZ-type CK production have not been identified yet.

CK biosynthesis is regulated by various internal and environmental factors (Hirose et al., 2008; Zwack and Rashotte, 2015). Nitrogen nutrition, which is available mostly as nitrate and ammonium in soil (Miller and Cramer, 2004; Kirk and Kronzucker, 2005; Xu et al., 2012; Kiba and Krapp, 2016; Coskun et al., 2017), is one of the major factors. Studies in *Arabidopsis* showed that *IPT3* and *CYP735As* are induced by a nitrate-specific signal to boost iP- and tZ-type CK accumulation in *Arabidopsis* (Miyawaki et al., 2004; Takei et al., 2004; Engelsberger and Schulze, 2012; Maeda et al., 2018). *OsIPT4* and *OsIPT5* are not induced by nitrate itself but by ammonium and glutamine with accompanying accumulation of iP-type and tZ-type CKs, indicating that *de novo* iP- and tZ-type CK biosynthesis is regulated by glutamine-related signals in rice (Kamada-Nobusada et al., 2013). A *CYP735A* gene in *Oryza longistaminata*, a wild rice, was shown to be upregulated by ammonium nitrate but the signal responsible for the upregulation is not known (Shibasaki et al., 2021). Thus, plants have dual systems, nitrate-specific and glutamine-related systems, to modulate the iP-type and tZ-type CK biosynthesis in response to nitrogen availability. Although the underlying mechanism is not understood, cZ-type CK levels are generally increased in response to biotic and abiotic stresses, such as phosphate starvation, low temperature, drought, and salt treatment (Vyroubalová et al., 2009; Macková et al., 2013; Vanková et al., 2014; Silva-Navas et al., 2019).

Previous studies have suggested that there is biological relevance to side-chain variations of CKs (Schmitz and Skoog, 1972; Mok et al., 1978; Werner et al., 2003; Spíchal et al., 2004; Gajdošová et al., 2011; Stolz et al., 2011; Choi et al., 2012; Osugi and Sakakibara, 2015; Schäfer et al., 2015). However, it is only recently that studies in *Arabidopsis* demonstrated it. The abundance of iP- and tZ-type CKs is higher than that of cZ-type (Osugi and Sakakibara, 2015; Osugi et al., 2017; Kiba et al., 2019) and tZ-type CKs are the most active, while cZ-type CKs are the least active in bioassays in *Arabidopsis* (Gajdošová et al., 2011; Kudo et al., 2012). *Arabidopsis* CK receptors have different affinities toward side-chain variants (Spíchal et al., 2004; Romanov et al., 2006; Stolz et al., 2011). For example, AHK2 and AHK4/WOL/CRE1 have high affinity to both iP and tZ, while AHK3 binds to iP with a ten times lower affinity than tZ in *in vitro* binding assays. Consistent with the bioassay, the affinity of AHKs to cZ is significantly lower compared with that of iP and tZ (Spíchal et al., 2004; Romanov et al., 2006; Stolz et al., 2011). Furthermore, characterization of the *Arabidopsis cyp735a1 cyp735a2* mutant that is impaired in tZ-type side-chain modification has unequivocally shown that the function of tZ-type CKs is shoot growth promotion, and it is different from iP-type and cZ-type CKs (Kiba et al., 2013). Thus, tZ-type CKs are the major active variants possessing a specific function in shoot growth, while cZ-type CKs are the weakly active or inactive forms with minor roles in *Arabidopsis*.

Although the occurrence of the three side-chain variants is ubiquitous among plants (Gajdošová et al., 2011), several lines of evidence suggest that their functional differentiation may not be as universal. For instance, the tZ-type CKs is not consistently more abundant than the cZ-type CKs among all plants. Plants that contain higher levels of cZ-type CKs than tZ-type can be found across the whole evolutionary tree of land plants (Gajdošová et al., 2011). Interestingly, many crops, including pea, potato, oats, maize and rice, are cZ-type-abundant plants (Gajdošová et al., 2011; Schäfer et al., 2015). Furthermore, cZ-type CKs are shown to be as active as tZ-type in some bioassays, including cytokinin-responsive gene expression assay in maize cultured cell (Yonekura-Sakakibara et al., 2004) and seminal root growth inhibition assay in rice (Kudo et al., 2012). Consistently, cytokinin receptors of maize (ZmHK1) and rice (OsHK3 and OsHK4) were shown to have similar affinity toward cZ and tZ (Lomin et al., 2011; Choi et al., 2012). However, the exact function of each of these variants in plants other than *Arabidopsis*, including monocot and cZ-type-abundant plants, is still unknown and remains to be determined.

Rice is a monocot plant possessing a significantly distinctive CK side-chain profile from *Arabidopsis*; cZ-type CKs comprise more than 80% of total CK in various tissues (Kojima et al., 2009; Kudo et al., 2012; Osugi and Sakakibara, 2015). In this study, we characterized the consequence of disruption of rice *CYP735A*s (*CYP735A3* and *CYP735A4*) and demonstrated that these genes are vital for tZ-type CK biosynthesis. Our results also revealed that tZ-type CK biosynthesis is regulated by phytohormones and dual nitrogen signals and that tZ-type CKs act to promote root and shoot growth in rice. These results suggest that the physiological role of tZ-type CKs in rice is different from that in *Arabidopsis* despite the fact that the biosynthetic mechanism is conserved. The function of tZ-type CKs might be optimized for each plant species depending on the body plan and/or growth environments.

## Results

### *CYP735A3* and *CYP735A4* complement the Arabidopsis *cyp735a1 cyp735a2* mutant

In *Oryza sativa* genome, *CYP735A3* and *CYP735A4* encode proteins with high amino acid sequence homology to CYP735A1 and CYP735A2, which are cytochrome P450 monooxygenases for tZ-type CK biosynthesis in *Arabidopsis* (Fig. S1) (Hansen et al., 2021). The *cyp735a1 cyp735a2* mutant (*atcypDM)* is deficient in tZ-type CKs resulting in shoot growth retardation (Kiba et al. 2013). To assess whether these genes encode enzymes that catalyze the same reaction as CYP735A1 and CYP735A2, we expressed them in *atcypDM* under the control of cauliflower mosaic virus (CaMV) 35S promoter. Accumulation of *CYP735A3* or *CYP735A4* transcripts was confirmed by quantitative RT-PCR analysis (qRT-PCR) in two independent lines of transgenic plants CYP735A3-ox or CYP735A4-ox, respectively (Fig. 1A). When grown on soil, all the transgenic lines developed a rosette with significantly greater diameter than *atcypDM* (Fig. 1B). Both *CYP735A3* and *CYP735A4* expression resulted in an increase of tZ-type CK (tZ and its conjugates) concentration with a concomitant decrease of iP-type CK (iP and its conjugates) concentration compared with *atcypDM*, but no significant differences in cZ-type CK (cZ and its conjugates) concentration were observed (Fig. 1C, Table S1). These results demonstrate that CYP735A3 and CYP735A4 catalyze the same reaction as CYP735A1 and CYP735A2, which is the *tran*s-hydroxylation of CK side chain to synthesize tZ-type CKs from iP-type CKs.

**Figure 1.**
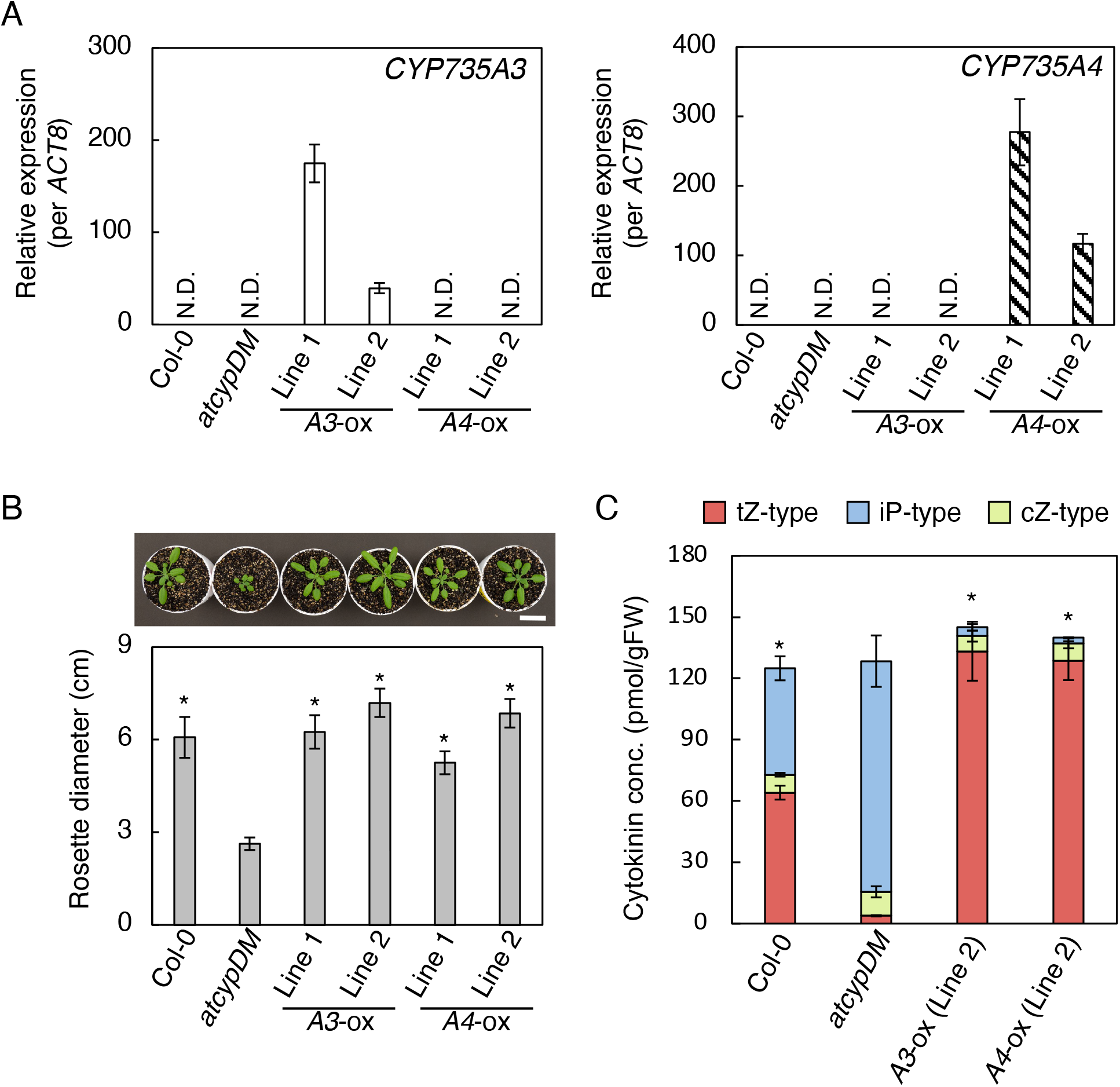
Recovery of *trans*-zeatin-type cytokinin deficiency in the *Arabidopsis* mutant *cyp735a1 cyp735a2* by *CYP735A3* or *CYP735A4*. (A) Quantitative RT-PCR analysis of *CYP735A3* (left) or *CYP735A4* (right) expression in whole seedlings of Col-0, *cyp735a1 cyp735a2* mutant (*atcypDM*), and transgenic plants constitutively expressing *CYP735A3* (*A3*-ox) or *CYP735A4* (*A4*-ox) in *atcypDM* background. Seedlings were grown on 1/2x MS agar plates for 12 days before harvest. Expression levels were normalized using *ACT8* as an internal control. Error bars represent standard deviation (n=4 independent pools of more than ten plants). N.D., under the quantification detection limit. (B) Rosette diameter of 28-day-old Col-0, *atcypDM*, and transgenic plants constitutively expressing *CYP735A3* (*A3*-ox) or *CYP735A4* (*A4*-ox) in *atcypDM* background grown on soil. A photograph of representative plants is shown. Error bars represent standard deviation (n=7). Asterisks indicate statistically significant differences compared with *atcypDM* (*p*<0.01, Student’s *t*-test). Scale bar, 3 cm. (C) Cytokinin concentrations in whole seedlings of Col-0, *atcypDM* and transgenic plants constitutively expressing *CYP735A3* (*A3*-ox) or *CYP735A4* (*A4*-ox) in *atcypDM* background. Seedlings were grown on 1/2x MS agar plates for 12 days before harvest. Error bars represent standard deviation (n=4 independent pools of more than ten plants). Asterisks indicate statistically significant differences in tZ-type cytokinin concentration compared with *atcypDM* (*p*<0.01, Student’s *t*-test). The concentration of each cytokinin molecular species in roots and shoots is shown in Table S1. gFW, gram fresh weight; tZ-type, *trans*-zeatin and its conjugates; cZ-type, *cis*-zeatin and its conjugates; iP-type, *N*^6^-(Δ^2^-isopentenyl)adenine and its conjugates.

### The spatial expression pattern of *CYP735A3* and *CYP735A4*

Expression levels of *CYP735A3* and *CYP735A4* were analyzed by qRT-PCR in various organs of rice in seedling to early reproductive phases (Fig. 2). At seedling phase, transcripts of *CYP735A*s were detected both in roots and shoots but they were more abundant in roots as reported previously (Tsai et al., 2012) (Fig. 2A). In later phases tested, the accumulation of the *CYP735A3* transcript was the highest around the vegetative shoot apex, while that of *CYP735A4* transcript was prominent in the leaf blade of plants at vegetative and early reproductive phases (Fig. 2B).

**Figure 2.**
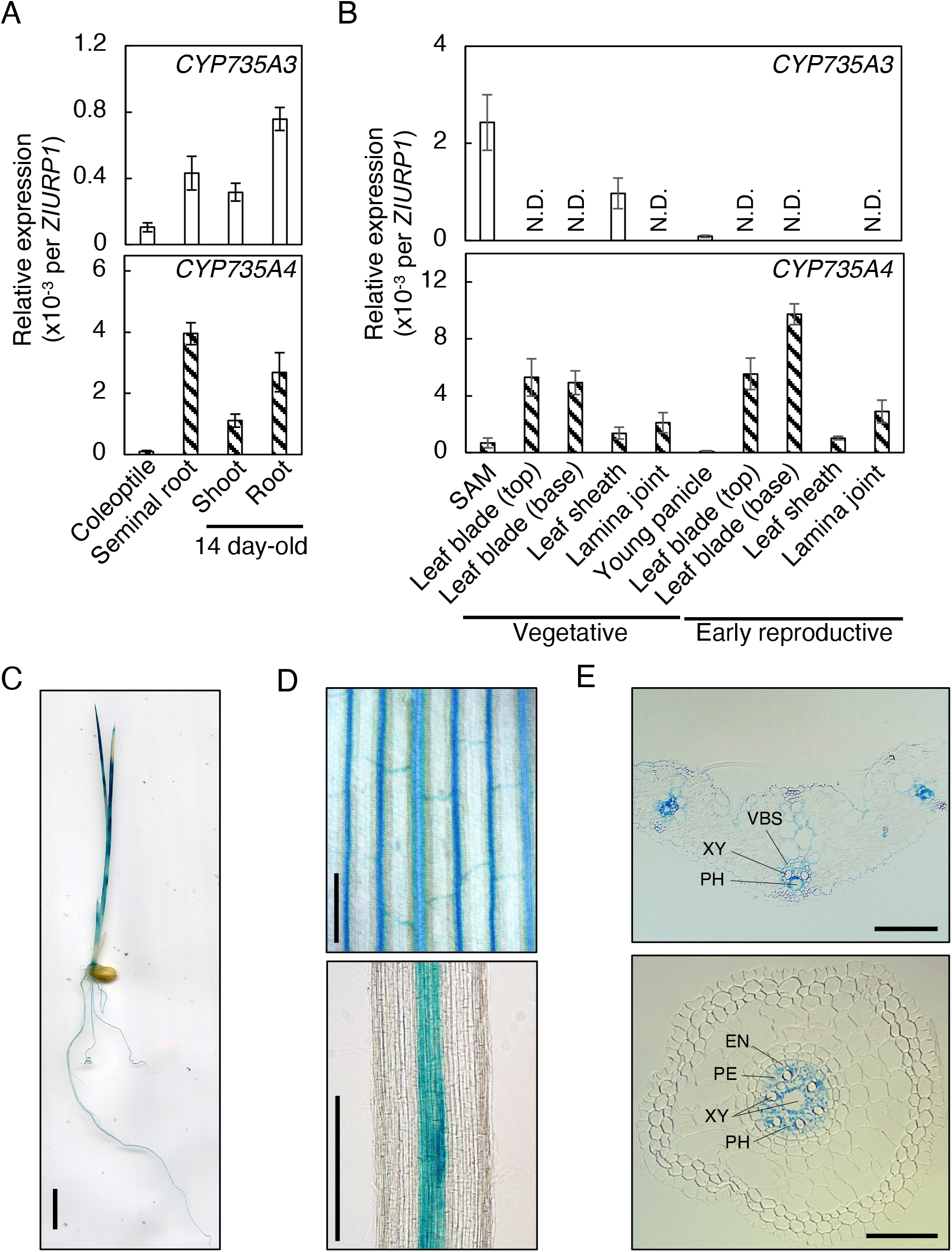
Expression pattern of *CYP735A3* and *CYP735A4* in various tissues. (A) Expression of *CYP735A3* and *CYP735A4* in Nipponbare seedlings. Coleoptiles and seminal roots, and roots and shoots were harvested 4 days and 14 days, respectively, after imbibition. Error bars represent standard deviation of biological replicates (n=4-5 independent pools of more than 2 plants). (B) Expression of *CYP735A3* and *CYP735A4* in Nipponbare plants in vegetative or early reproductive stages. 42-day-old (vegetative stage) and older plants (reproductive stage) were used. The shoot apex parts of 42-day-old plants were collected as shoot apical meristem (SAM). Leaf blade, leaf sheath and lamina joint were harvested from the youngest fully expanded leaf. The young panicle is a mixture of 2 mm to 21 mm long young panicles collected from plants at the reproductive stage. Error bars represent standard deviation of biological replicates (n=4 except for SAM and young panicle, n=4 and n=7 independent pools of more than 2 plants for SAM and young panicle, respectively). N.D., under the quantification detection limit. (C-E) Representative images of GUS staining obtained from *proCYP735A3:GUS* transgenic seedlings grown for 8 days after imbibition. Pictures of a whole seedling (C), first leaf (D, upper), crown root (D, lower), cross section of the first leaf (E, upper), and cross section of seminal root (E, lower) are presented. VBS, vascular bundle sheath; XY, xylem; PH, phloem; EN, endodermis; PE, pericycle. Scale bars, (C) 1 cm; (D) 200 μm; (E) 50 μm.

In order to localize *CYP735A*s expression at the tissue level, we used GUS staining technique. For this purpose, we generated transgenic plants harboring a fusion of the 3.6 kb *CYP735A* upstream sequence and the *GUS* reporter gene (proCYP735A3:GUS and proCYP735A4:GUS). Similar staining patterns were observed in multiple independent T1 lines of proCYP735A3:GUS (Figs. 2C-E). However, no staining was detected in any of the proCYP735A4:GUS lines (data not shown), suggesting that the 3.6 kb upstream sequence of *CYP735A4* was not sufficient to drive expression. Consistent with the qRT-PCR results (Fig. 2A), staining was detected both in roots and shoots of proCYP735A3:GUS seedlings (Fig. 2C). The GUS activity was restricted to the vascular bundle (Fig. 2D) and observation of cross-sections revealed that the staining was pronounced in vascular parenchyma cells in the shoot, and vascular parenchyma and pericycle cells in the root (Fig. 2E).

### *CYP735A3* and *CYP735A4* play a central role in tZ-type cytokinin biosynthesis in rice

To examine the role of *CYP735A*s in CK side-chain modification in rice, we disrupted *CYP735A3* and *CYP735A4* by CRISPR-Cas9 system. Two transfer RNA-based-multiplex CRISPR-Cas9 vectors, each harboring three different guides targeting unique sequences of *CYP735A3* and *CYP735A4*, were used to edit *CYP735A3* and *CYP735A4* (Figs. S2 and S3). Two independent lines with mutations in both *CYP735A3* and *CYP735A4* were identified. In a T1 line, the *cyp735a3-1* and *cyp735a4-1* alleles were found as homozygote and heterozygote, respectively (Fig. S2). In another T1 line, the *cyp735a3-2* and *cyp735a4-2* alleles were found as heterozygote and homozygote, respectively (Fig. S3). The *cyp735a3-1* single, *cyp735a4-2* single, *cyp735a3-1 cyp735a4-1* (*a3a4-1*) double and *cyp735a3-2 cyp735a4-2* (*a3a4-2*) double mutants were isolated from progenies of these lines. The *cyp735a3-1* and *cyp735a3-2* are one-base insertions that shift the reading frame and yield predicted proteins with no similarity to any Non-redundant UniProtKB/SwissProt sequences and without the cytochrome P450 domain (Fig. S4). The *cyp735a4-1* and *cyp735a4-2* mutants are 263- and 298-base deletions, respectively. The 263- and 298-base deletions resulted in a predicted protein lacking “A helix”, a secondary structure conserved among cytochrome P450s (Midlik et al., 2021), and an in-frame stop codon at 4 codons after the mutation, respectively (Fig. S5). Since the proteins predicted to be produced in all the mutants (except for *cyp735a4-1*) are apparently inactive as cytochrome P450, we concluded that *cyp735a3-1, cyp735a3-2*, and *cyp735a4-2* are null alleles.

CK concentrations in roots and shoots were measured in wild type (WT) strain Nipponbare (NB), vector control (VC), and mutant strains *cyp735a3-1, cyp735a4-2, a3a4-1* and *a3a4-2* seedlings grown in hydroponic culture (Fig. 3, Tables S2 and S3). The concentrations of tZ-type CK and iP-type CK were comparable to WT in *cyp735a3-1* (Fig. 3A). In contrast, *cyp735a4-2* had slightly decreased tZ-type CK and slightly increased iP-type CK concentrations in roots and shoots as compared to the WT (Fig. 3B). In *a3a4-1* and *a3a4-2*, the concentration of tZ-type CKs reduced dramatically to 3.5 to 8.8% of the wild-type levels in roots and shoots, while that of iP-type CK increased more than 2 -fold. In the root of double mutants, iP-type CK levels were increased so dramatically that the loss of tZ-type CK was compensated in terms of the sum of iP- and tZ-type CKs (iP-/tZ-type CK quantity). On the other hand, in the shoot, iP-type CK levels were increased but the iP-/tZ-type CK quantity was significantly lower compared with WT (Fig. 3; Tables S2 and S3). The cZ-type CK levels were not altered in any of the mutants (Fig. 3, Tables S2 and S3). Since the CK profiles of *a3a4-1* and *a3a4-2* are distorted in a similar manner, *cyp735a4-1* seems to be a null allele. These results show that tZ-type CKs are mostly synthesized by CYP735A3 and CYP735A4 in rice.

**Figure 3.**
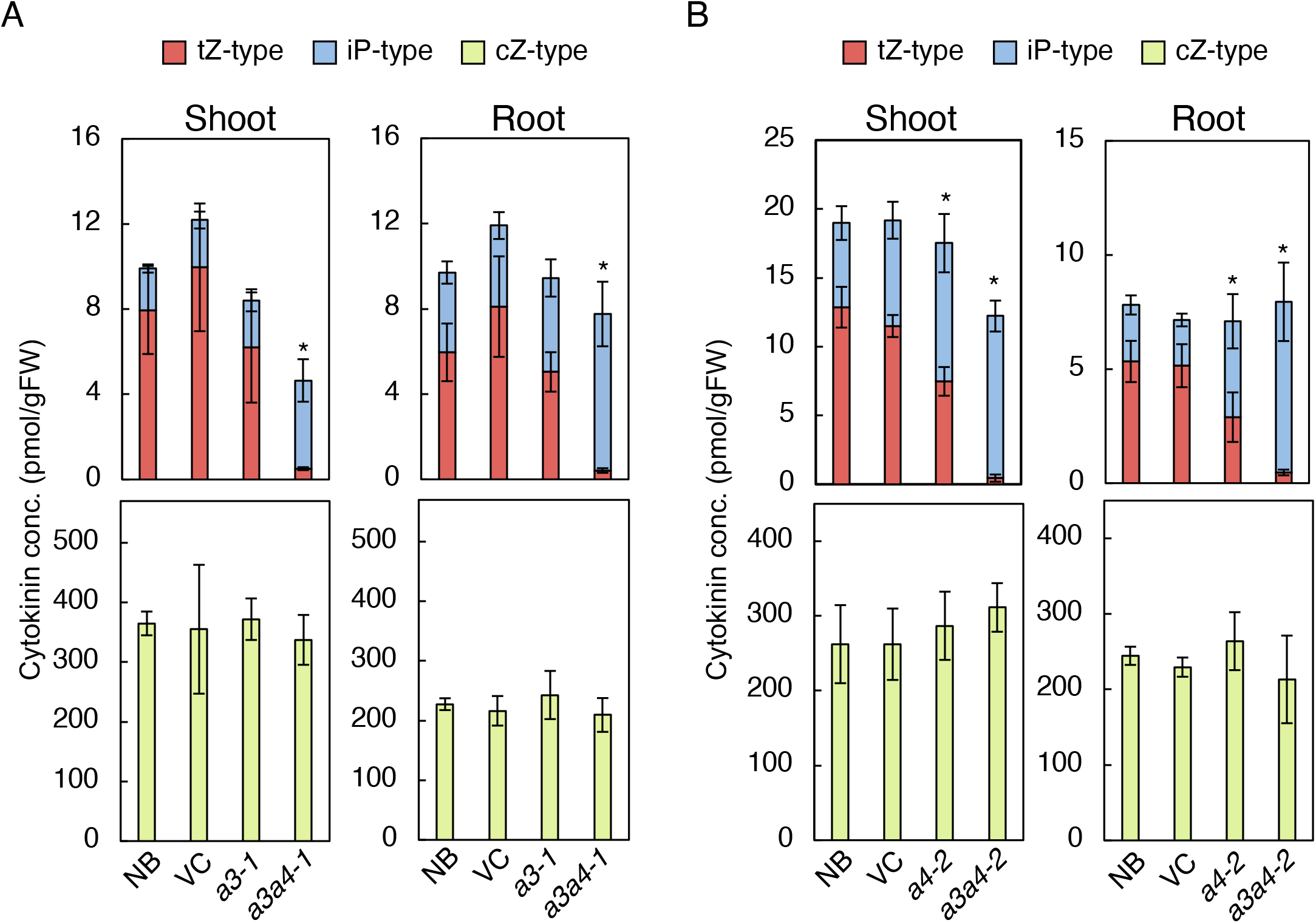
The *cyp735a3 cyp735a4* double mutants show deficiency in *trans*-zeatin-type cytokinins. Cytokinin concentrations in roots and shoots of WT strain Nipponbare (NB), vector control (VC), and mutant strains *cyp735a3-1* (*a3-1*), *cyp735a4-2* (*a4-2*), *cyp735a3-1 cyp735a4-1* (*a3a4-1*), and *cyp735a3-2 cyp735a4-2* (*a3a4-2*). Seedlings were grown hydroponically for 14 days (A) or 12 days (B) before shoots and roots were harvested. Error bars represent standard deviation of biological replicates (n=3-6 independent pools of more than ten plants for A, n=6-8 independent pools of more than ten plants for B). Asterisks indicate statistically significant differences in tZ-type cytokinin concentration compared with NB (*p*<0.05, Student’s *t*-test). The concentration of each cytokinin molecular species in roots and shoots is shown in Table S2 and Table S3. gFW, gram fresh weight; tZ-type, *trans*-zeatin and its conjugates; cZ-type, *cis*-zeatin and its conjugates; iP-type, *N*^6^-(Δ^2^-isopentenyl)adenine and its conjugates.

### Disruption of *CYP735A3* and *CYP735A4* retards growth of rice

To explore the physiological role of tZ-type CKs in rice, we examined the growth and development of *cyp735a* single and double mutants from seedling phase to maturation phases. Although *cyp735a3-1* and *cyp735a4-2* single mutants were indistinguishable from control plants (WT and VC) on our growth conditions, similar significant alterations in growth and development were observed in *a3a4-1* and *a3a4-2* double mutants (Figs. 4, 5 and S6).

**Figure 4.**
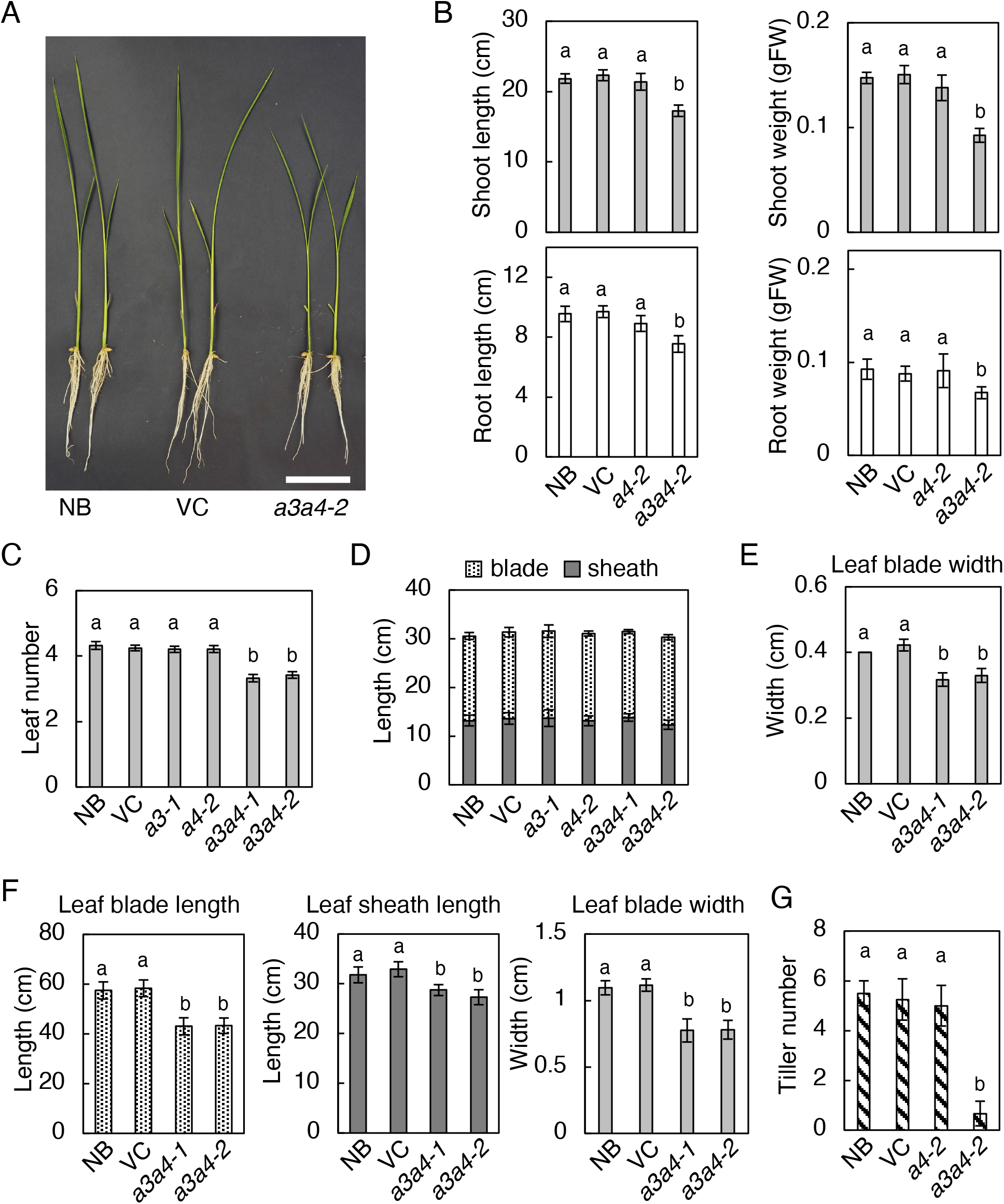
Disruption of *CYP735A*s alters rice growth in seedling to vegetative phases. (A) A representative image of Nipponbare (NB), vector control (VC), and *cyp735a3-2 cyp735a4-2* (*a3a4-2*) seedlings grown 12 days in hydroponic culture. Scale bar, 5 cm. (B) Quantification of shoot length, shoot fresh weight, root length, and root fresh weight in 12-day-old seedlings. NB, VC, *cyp735a4-2* (*a4-2*), and *a3a4-2* seedlings were grown in hydroponic culture. (C) Leaf number at 15 days after germination (D, E) Leaf sheath and leaf blade length (D) and leaf width (E) of the fully expanded third leaf of NB, VC, *cyp735a3-1* (*a3-1*), *cyp735a4-2* (*a4-2*), *cyp735a3-1 cyp735a4-1* (*a3a4-1*), and *a3a4-2* seedlings grown on soil. (F) Quantification of leaf blade length, leaf sheath length, and leaf width of the fully expanded ninth leaf in NB, VC, *a3a4-1*, and *a3a4-2* grown on soil. (G) Tiller number of NB, VC, *a3a4-1*, and *a3a4-2* grown on soil for 35 days. Error bars represent standard deviation of biological replicates (B, n=6-8; C, n=8-12; D, n=8-10; E, n=8-10; F, n=8-12; G, n=4-8). Different lowercase letters indicate statistically significant differences as indicated by Tukey’s HSD test (*p*<0.05).

**Figure 5.**
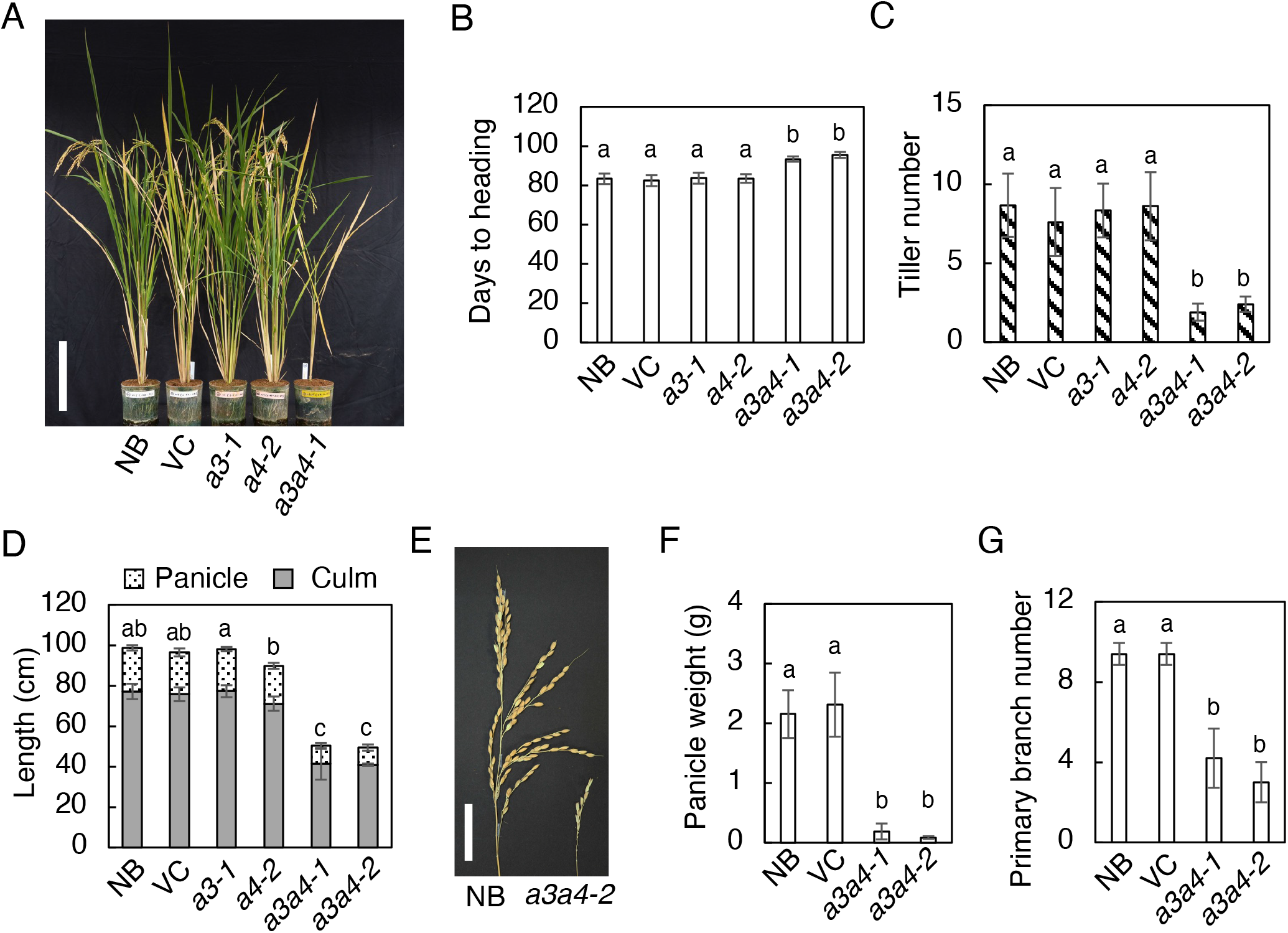
Disruption of *CYP735A*s alters rice growth in reproductive to maturation phases. (A) A representative image of Nipponbare (NB), vector control (VC), and *cyp735a3-1* (*a3-1*), *cyp735a4-1* (*a4-2*), and *cyp735a3-1 cyp735a4-1* (*a3a4-2*) plants grown 15 weeks on soil. Scale bar, 20 cm. (B-D) Heading date (B), panicle and culm length (C), and tiller number (D) of NB, VC, *a3-1, a4-2, cyp735a3-1 cyp735a4-1* (*a3a4-1*), and *a3a4-2*. (E) A representative picture of mature panicles from NB and *a3a4-2*. Scale bar, 5 cm. (F, G) Panicle weight on the main stem (F) and number of primary branches per panicle (G). Error bars represent standard deviation of biological replicates (B, n=3-12; C, n=5-12; D, n=9-12; F, n=5-10; G, n=5-10). Different lowercase letters indicate statistically significant differences as indicated by Tukey’s HSD test (*p*<0.05).

The double mutant seedlings showed a decrease in the length and fresh weight of the shoot and root compared with control (Figs. 4A, 4B, S6A, S6B). To determine the cause of the decrease in shoot length and fresh weight, leaf number and leaf size were measured (Figs. 4C-E). Leaf number of double mutant seedlings at 15 days after germination was reduced compared to control (Fig. 4C), indicating that the rate of leaf formation is slower in the double mutant. Leaf blade and sheath length of fully expanded third leaf was similar for the double mutant and control (Fig. 4D), while leaf blade width was decreased in the double mutant (Fig. 4E). These results indicated that a decrease in leaf number accounts for the reduction in shoot length and a decrease in leaf number and leaf width contributes to the reduction in shoot fresh weight. Reduced shoot length phenotype of the double mutant continued into later vegetative growth stage (Figs. S6C and S6D), but it cannot be attributed only to a decrease in leaf number because leaf size, namely blade length, blade width and sheath length, was significantly reduced at this stage as observed in fully expanded ninth leaf (Fig. 4F). In addition, the double mutant developed significantly fewer tillers compared with control (Figs. 4G and 5C), suggesting that the activities of shoot apical and axillary meristems are impaired.

All control plants headed about 83 days after germination (n=19) but double mutant plants either died without heading from the main stem (16 out of 24 plants) or showed delayed heading (8 out of 24 plants) (Figs. 5A and 5B). Panicle length, panicle weight and number of primary panicle branch were all significantly reduced in the double mutant compared to the control (Figs. 5D-G), suggesting that inflorescence meristem activity is reduced. The double mutant set few but viable seeds, indicating that reproductive organ development is not severely affected. Together, these results suggest that the tZ-type CKs regulate both root and shoot growth, most possibly through controlling meristem activity.

### The cytokinin activity is reduced both in roots and shoots of the *cyp735a3 cyp735a4* mutant

The growth defects observed in *a3a4* double mutant are reminiscent of those observed in rice mutants with diminished CK activities such as type-A *RESPONSE REGULATOR* (*RR*) *OsRR6* overexpressing lines (Hirose et al., 2007), type-B *RR* triple mutant *rr21/22/23* (Worthen et al., 2019) and cytokinin receptor double mutant *hk5 hk6* (Burr et al., 2021). To examine whether the CK activity is reduced in the double mutant, we analyzed the expression levels of immediate-early CK-inducible type-A *RR*s by qRT-PCR (Fig. 6). The expression level of all tested type-A *RR*s, except for that of *OsRR1* in roots, was significantly lower in the mutants compared with WT in both roots and shoots (Figs. 6A and 6B). On the other hand, the expression of non-CK-inducible type-B *RR*s, *OsRR21* and *OsRR23*, was not reduced (Figs. 6C and 6D). Thus, cytokinin activity in roots and shoots of *a3a4* is diminished, suggesting that the *a3a4* growth phenotype is caused by decreased CK activity.

**Figure 6.**
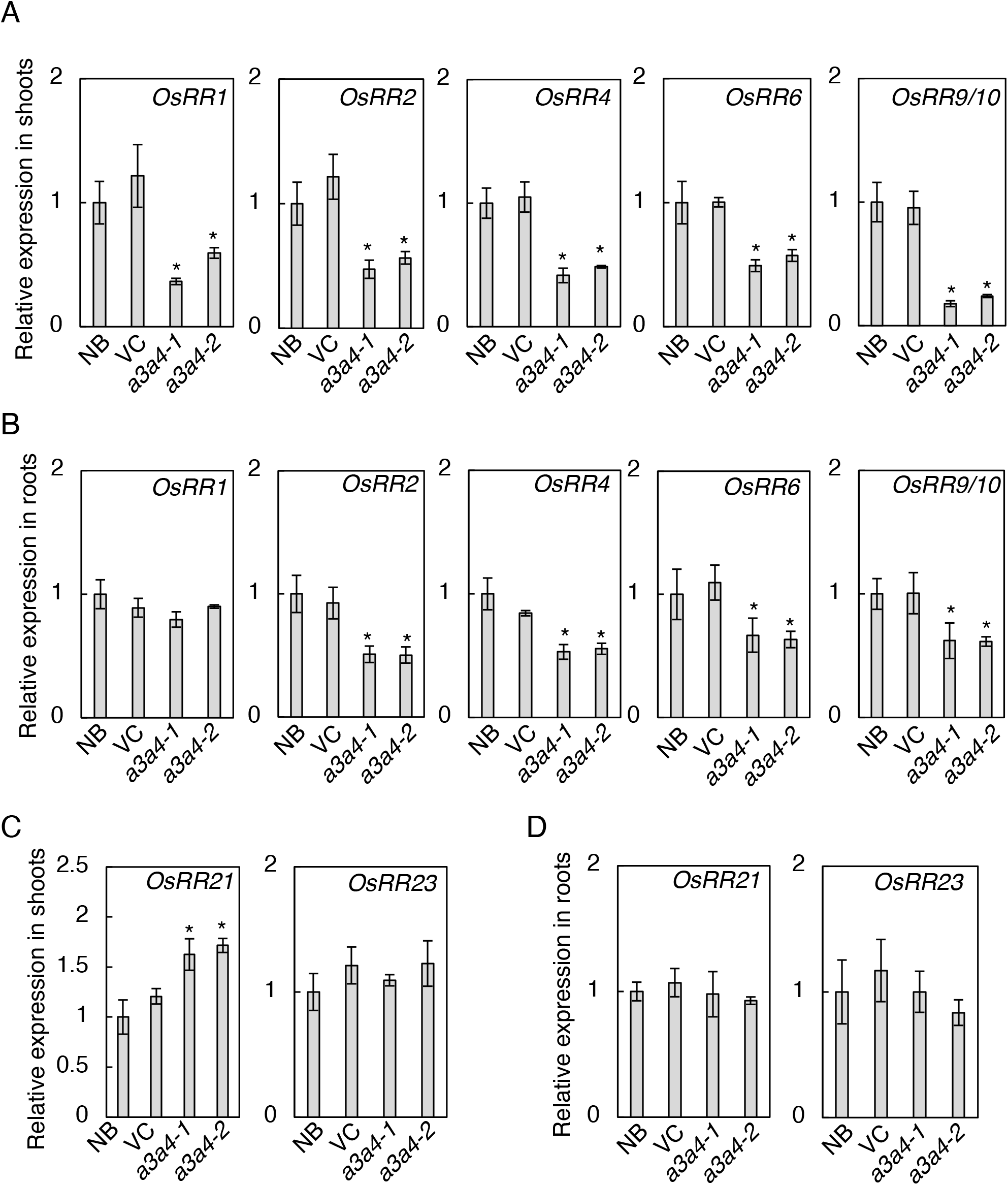
Expression of cytokinin inducible type-A response regulator genes is altered in the *cyp735a3 cyp735a4* mutant. (A, B) Transcript levels of cytokinin inducible type-A response regulator genes, *OsRR1, OsRR2, OsRR*4, *OsRR6*, and *OsRR9/10*, in shoots (A) and roots (B) of Nipponbare (NB), vector control (VC), *cyp735a3-1 cyp735a4-1* (*a3a4-1*) and *cyp735a3-2 cyp735a4-2* (*a3a4-2*) seedlings. *OsRR9/10* indicates that the primer set used could not distinguish *OsRR9* and *OsRR10*. (C, D) Transcript levels of type-B response regulator genes, *OsRR21* and *OsRR23*, in the shoot. (C) and in the root (D) of NB, VC, *a3a4-1* and *a3a4-2* seedlings. Seedlings were grown hydroponically for 10 days before roots and shoots were harvested. Expression levels were quantified by qRT-PCR analysis, normalized to *ZIURP1* as an internal control, and are shown relative to the value of NB. Error bars represent standard deviation of biological replicates (n=4 independent pools of 2 plants). Asterisks indicate statistically significant differences compared to NB (*p*<0.05, Student’s *t*-test).

### Regulation of *CYP735A3* and *CYP735A4* expression by phytohormones

To obtain insights into the physiological relevance of tZ-type CK biosynthesis in rice, we analyzed the responses of *CYP735A*s to phytohormones. In *Arabidopsis*, the expression of *CYP735A1* and *CYP735A2* was shown to be upregulated by CK and downregulated by auxin and abscisic acid (Takei et al., 2004). Tsai et al (2012) analyzed the effect of these phytohormones on *CYP735A3* and *CYP735A4* expression by the NanoString nCounter system but could not detect *CYP735A3* under some conditions using this system. To fully characterize the response of *CYP735A*s to these phytohormones in rice, we analyzed their expression in roots by qRT-PCR. Nipponbare seedings treated with phytohormones were used for the analysis. *OsRR4*/*OsRR6* (Kudo et al., 2012; Tsai et al., 2012), *OsSIPP2C1* (Li et al., 2013) and *OsGH3*.*2* (Du et al., 2012) were used as positive controls for CK, IAA and ABA responses, respectively. Consistent with the NanoString analysis (Tsai et al., 2012), we found that the *CYP735A4* transcript was decreased by CK (iP and tZ), IAA and ABA (Fig. 7). Our qRT-PCR analysis showed that *CYP735A3* is also downregulated by CK, IAA and ABA (Fig. 7). Thus, rice *CYP735A*s and *Arabidopsis CYP735A*s are regulated similarly by IAA and ABA, but differently by CK.

**Figure 7.**
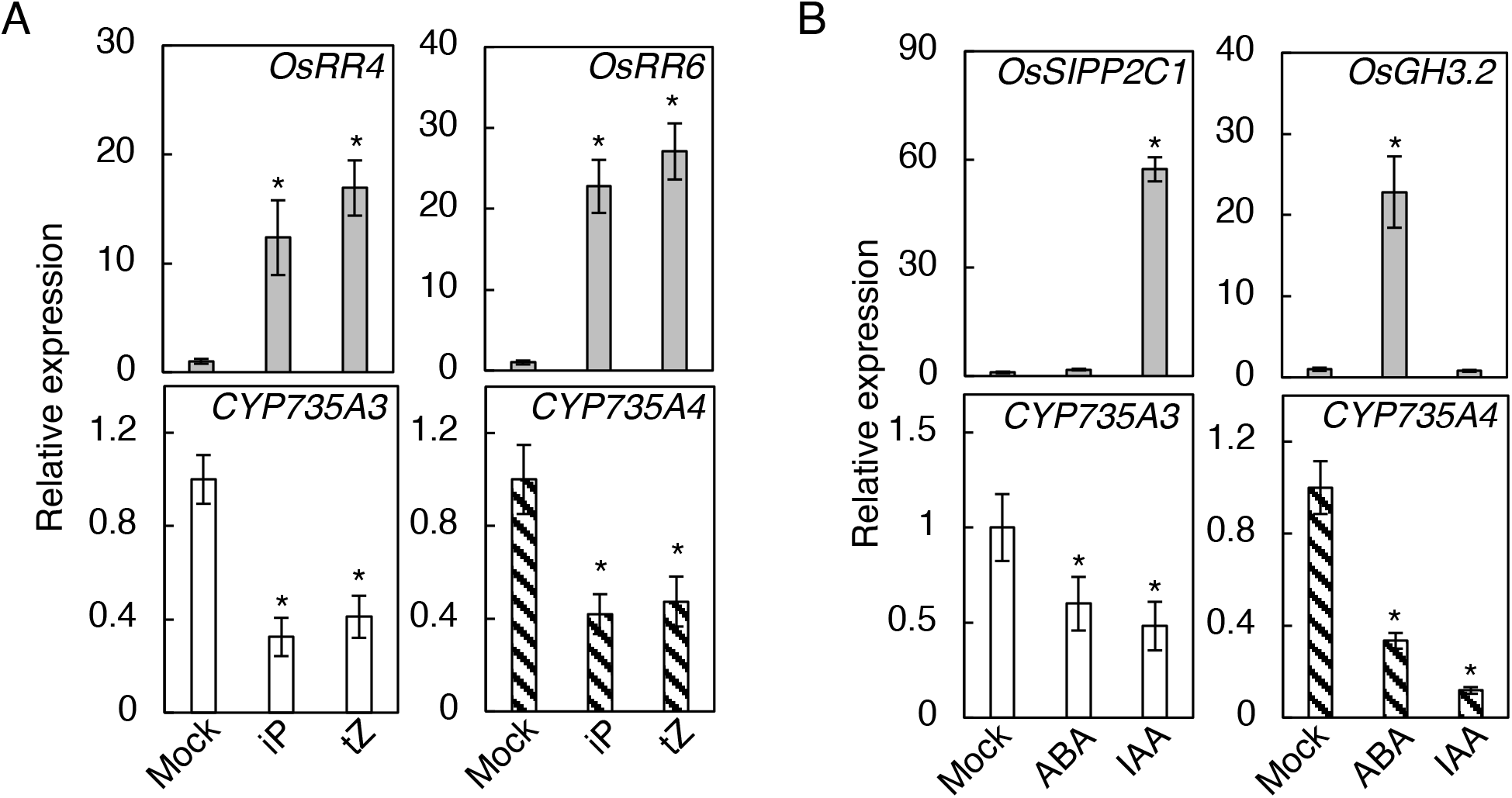
Expression of *CYP735A3* and *CYP735A4* in response to phytohormones. (A) Expression of *CYP735A*s in response to cytokinin. Nipponbare (NB) seedlings grown for 12 days were treated with 0.001% DMSO (Mock), 1 μM iP (iP), or 1 μM tZ (tZ) and roots were harvested after 2 h. (B) Expression of *CYP735A*s in response to abscisic acid (ABA) and auxin. NB seedlings grown for 12 days were treated with 0.05% DMSO (Mock), 10 μM indole-3-acetic acid (IAA) or 50 μM ABA and roots were harvested after 2 h. Expression levels were quantified by qRT-PCR analysis, normalized to *ZIURP1* as an internal control, and are shown relative to the value in mock treatment. Asterisks indicate statistically significant differences compared with Mock (*p*<0.05, Student’s *t*-test). Error bars represent standard deviation (n=4 independent pools of more than 2 plants).

### Regulation of *CYP735A3* and *CYP735A4* expression by nitrogen sources

Nitrogen nutrition is one of the major environmental factors regulating CK biosynthesis, including tZ-type CK biosynthesis, for plant growth optimization (Takei et al., 2001b; Kamada-Nobusada et al., 2013; Ohashi et al., 2017; Osugi et al., 2017; Maeda et al., 2018; Sakakibara, 2021; Shibasaki et al., 2021; Kawai et al., 2022). In *Arabidopsis, CYP735As* are upregulated by the nitrate-specific signal and play a crucial role in nitrate-induced accumulation of tZ-type CKs (Engelsberger and Schulze, 2012; Maeda et al., 2018). To test whether *CYP735A*s of rice are nitrogen-inducible, we analyzed the expression levels of *CYP735A*s in roots supplemented with ammonium or nitrate (Fig. 8). *OsIPT4* and *OsNIA1* were used as indicators for glutamine-related and nitrate-specific responses, respectively (Kamada-Nobusada et al., 2013). *CYP735A3* was induced by both ammonium and nitrate in a manner similar to *OsIPT4*, while the expression of *CYP735A4* was upregulated only by nitrate like *OsNIA1* (Fig. 8A). To clarify whether *CYP735A*s are regulated by glutamine-related and/or nitrate-specific signal, we examined the effect of methionine sulfoximine (MSX), an inhibitor of glutamine synthetase (Fig. 8B) (Lam et al., 1996). MSX pre-treatment inhibited the nitrate-triggered induction of *CYP735A3* but did not affect *CYP735A4*. On the other hand, *CYP735A3* was upregulated by glutamine supplement in the presence of MSX, while *CYP735A4* was not. These responses of *CYP735A3* and *CYP735A4* to nitrogen sources and MSX are similar to those of *OsIPT4* and *OsNIA1*, respectively, indicating that *CYP735A3* and *CYP735A4* are regulated by glutamine-related and nitrate-specific signals, respectively. Together, these results suggested that tZ-type CK biosynthesis in rice is controlled by dual nitrogen signals, which is different from that in *Arabidopsis*.

**Figure 8.**
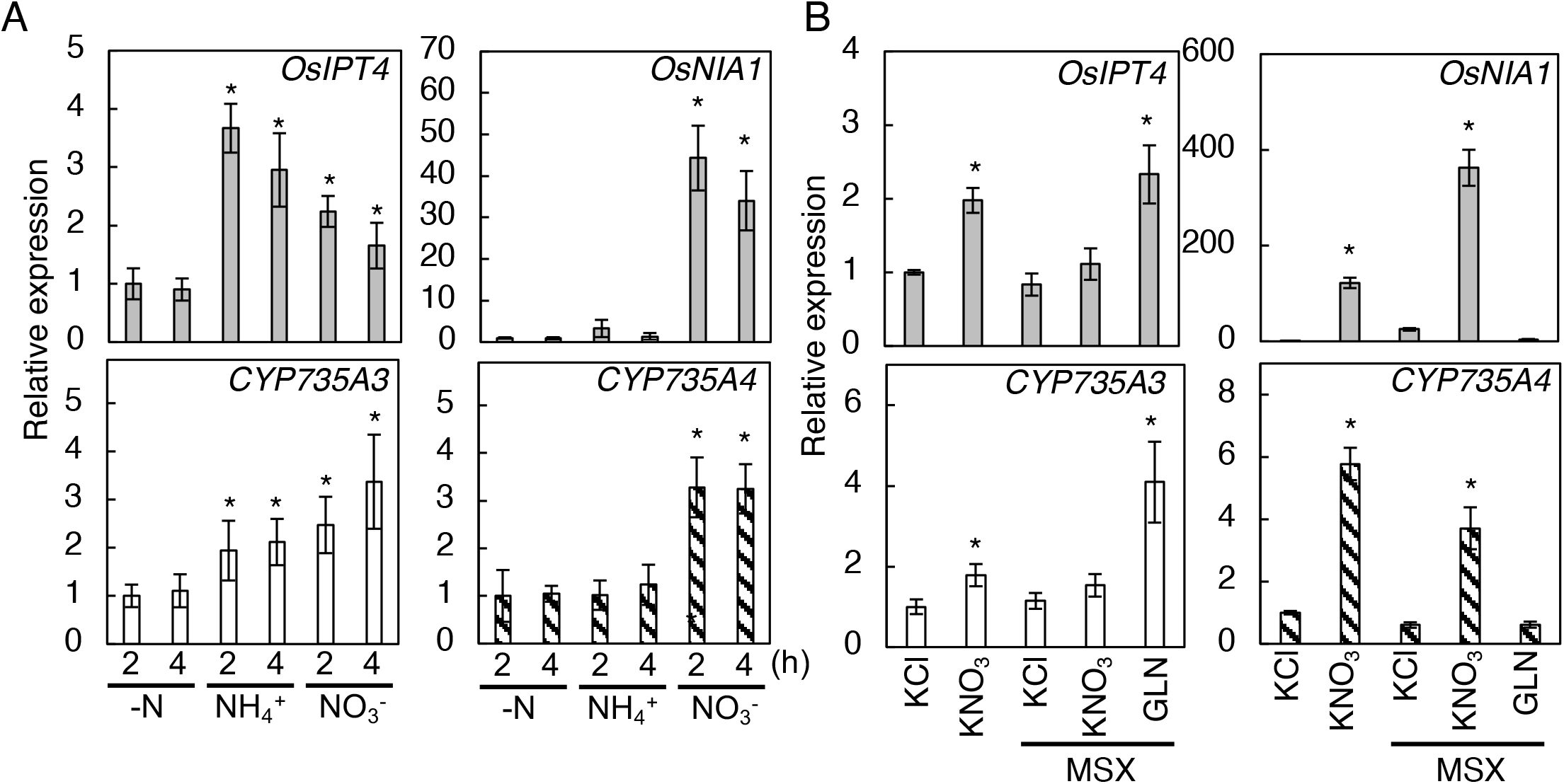
Expression of *CYP735A3* and *CYP735A4* in response to nitrogen sources. (A) Expression of *CYP735A*s in response to ammonium or nitrate supplement. Twelve-day-old NB seedlings hydroponically grown in nitrogen-free nutrient solution were incubated with 5 mM KCl (-N), 5 mM NH_4_Cl (NH_4_^+^), or 5 mM KNO_3_ (NO_3_^-^) for 2 h or 4 h. Expression levels were quantified by qRT-PCR analysis, normalized to *ZIURP1* as an internal control, and are shown relative to the value in 2 h of -N treatment. Error bars represent standard deviation (n=4 independent pools of 3 plants). (B) Effect of methionine sulfoximine on the induction of *CYP735A*s by nitrogen. Twelve-day-old NB seedlings hydroponically grown in nitrogen-free nutrient solution were pre-treated with 2 mM MSX for 2 h and then incubated with 5 mM KCl, 5 mM KNO_3_ or 50 mM glutamine (GLN) for 2 h. Asterisks indicate statistically significant differences compared with KCl or KCl+MSX (*p*<0.05, Student’s *t*-test). Expression levels were quantified by qRT-PCR analysis, normalized to *ZIURP1* as an internal control, and are shown relative to the value in KCl treatment. Error bars represent standard deviation (n=4 independent pools of 3 plants).

## Discussion

Angiosperms can be divided into four main categories in terms of their CK side-chain profile and number of cotyledons; iP/tZ-type-abundant dicot, cZ-type-abundant dicot, iP/tZ-type-abundant monocot and cZ-type-abundant monocot (Gajdošová et al., 2011). The function of some of these CKs has been demonstrated in a few dicotyledonous plant species. For example, in *Arabidopsis*, an iP/tZ-type-abundant dicot plant, CYP735As catalyze the biosynthesis of tZ-type CKs, which plays a specific role in shoot growth promotion (Kiba et al., 2013). Recently, it was suggested that CYP735A plays a similar role in *Jatropha curcas*, which is also an iP/tZ-type-abundant dicot plant (Cai et al., 2018). However, the function of CYP735A and tZ-type CKs in plants in other categories remains unknown. Here, we investigated the biosynthetic mechanism and function of tZ-type CKs in cZ-type-abundant monocot plant rice. Characterization of the *a3a4* double mutant demonstrated that *CYP735A*s play a central role in tZ-type CK biosynthesis in rice as in the iP/tZ-type-abundant dicot plants but that the physiological function of tZ-type CKs in rice is not the same as that in *Arabidopsis*, providing new insights into the biological significance of side-chain variations in plants.

*Arabidopsis* CYP735As catalyze the *trans*-hydroxylation of iP-type CKs to form tZ-type CKs (Takei et al., 2004; Kiba et al., 2013). Complementation test of *atcypDM* and CK quantification of *a3a4* demonstrated that CYP735A3 and CYP735A4 are the functional orthologs of *Arabidopsis* CYP735As (Figs. 1 and 3; Tables S1 to S3). The tZ-type CKs in *a3a4* was scarce but it did not completely disappear (Fig. 3; Tables S2 and S3). This is similar to what has been observed in the CYP735A-null mutants (*atcypDM*) of *Arabidopsis* and *J. curcas* (Kiba et al., 2013; Cai et al., 2018), suggesting the existence of minor alternative pathway(s) in both iP/tZ-type-abundant and cZ-type-abundant plants. The isomerization of cZ-type CKs is a possible pathway because *cis*-to-*trans* isomerization activity has been detected in various plants, including rice (Bassil et al., 1993; Gajdošová et al., 2011; Kudo et al., 2012). Although the isomerization pathway might be important under certain biological contexts, given that the pathway is minor even in the cZ-type-abundant rice, and that *CYP735A* homologs are found in all angiosperms whose genome is completely sequenced (Hansen et al., 2021), the CYP735A pathway seems to be the predominant pathway for tZ-type CK biosynthesis in angiosperms.

The *a3a4* mutants displayed severe growth retardation in vegetative and maturation phases (Figs 4, 5, and S5). We concluded that this phenotype is the result of diminished CK activity caused by a reduction in tZ-type CKs based on the following reasons: (i) the double mutant had severely reduced level of tZ-type CKs (Fig. 3; Tables S2 and S3); (ii) the cytokinin activity, measured by expression levels of immediate-early CK-inducible *RR* genes, was reduced in the double mutant (Fig. 6); (iii) The phenotypes of the double mutant is partly similar to those observed in rice mutants with severely reduced CK activity, such as *OsRR6* overexpressors (Hirose et al., 2007), *rr21/22/23* (Worthen et al., 2019) and *hk5 hk6* (Burr et al., 2021); (iv) our results were similar to the positive role of tZ-type CKs in rice growth suggested through the analysis of OsABCG18, a transporter involved in root-to-shoot CK translocation (Zhao et al., 2019); (v) similar conclusion was drawn from the data obtained from equivalent experiments on *CYP735A1* and *CYP735A2* in *Arabidopsis* (Kiba et al., 2013).

The cZ-type CKs are the dominant variants in terms of quantity comprising more than 80% of total CK in various rice tissues and are suggested to be an active CK that plays a role in normal growth and development in rice (Kojima et al., 2009; Choi et al., 2012; Kudo et al., 2012; Osugi and Sakakibara, 2015). Our CK quantification results showed that the levels of any of the cZ-type CK derivatives were not consistently altered in *a3a4* (Fig 3; Tables S2 and S3), indicating that cZ-type CKs are not directly relevant to the *a3a4* phenotype. The iP-type CKs are also considered to be an important variant in rice because of the identification of a receptor (OsHK6) with preferential affinity for iP (Choi et al., 2012). In *a3a4*, the iP-type CK levels were increased to compensate for the loss of tZ-type CK in the root but not sufficiently increased in the shoot (Fig. 3; Tables S2 and S3). From these results it could be interpreted that the cause of the *a3a4* phenotype is an alteration in CK side-chain profile (CK quality) but not CK quantity in the root and alteration in both CK quality and/or quantity in the shoot. Together, our results suggest that the function of tZ-type CKs is different from that of iP-type and cZ-type CKs at least in rice roots.

The phenotypes of *a3a4* were observed both in roots and shoots, including reduced root proliferation, delayed plastochron and decreased leaf size, panicle size and tiller number (Figs. 4, 5 and S5), indicating the extensive role of tZ-type CKs as growth enhancers in various organs in rice. Although it is difficult to simply compare the phenotypes of *a3a4* with those of *atcypDM* because monocot and dicot have remarkably different body plan, the major difference lies in the existence of phenotype in the root. The *a3a4* displayed reduced cytokinin activity and proliferation in the root (Figs. 4, 6 and S5), while no apparent difference was detected in *atcypDM* root (Kiba et al., 2013), suggesting that the role of tZ-type CKs in rice varies from that in *Arabidopsis*. The shoot specific-function of tZ-type CKs in *Arabidopsis* is partly explained by the ligand preference of cytokinin receptors (Spíchal et al., 2004; Romanov et al., 2006; Stolz et al., 2011; Kiba et al., 2013). Among the five genes encoding cytokinin receptors (OsHK3-6, CHARK) in rice, so far, the ligand preference of OsHK3, OsHK4 and OsHK6 has been investigated (Choi et al., 2012). However, no receptor with preferential affinity for tZ-type CKs has been identified and the mechanism underlying the functional specificity of tZ-type CKs in rice remains to be elucidated.

Consistent with the functional distinction of tZ-type CKs between rice and *Arabidopsis*, we also showed differences in responses to some internal and external cues of *CYP735A*s (Figs. 2, 7 and 8). Rice *CYP735A*s are repressed by CKs (Tsai et al. 2012), while *Arabidopsis CYP735A*s are induced (Takei et al., 2004), suggesting that rice and *Arabidopsis* have negative feedback and positive feedback regulation, respectively, on tZ-type CK function. The expression level of *CYP735A4* is high in leaves (Fig. 2) but that of *Arabidopsis CYP735A*s is not (Takei et al., 2004; Kiba et al., 2013). Responses of these genes to nitrogen supplement were also found to be different. *CYP735A3* and *CYP735A4* are upregulated by glutamine-related and nitrate-specific signals, respectively, in rice (Fig. 8) while *CYP735A*s of *Arabidopsis* are induced only by nitrate-specific signal (Engelsberger and Schulze, 2012; Maeda et al., 2018). Although ammonium is known to be the major form of inorganic nitrogen source in paddy field, rice utilizes not only ammonium but also nitrate generated by nitrification on the root surface as inorganic nitrogen sources (Kirk and Kronzucker, 2005; Yan et al., 2011). For *Arabidopsis*, which grows on aerobic soil, nitrate is the major inorganic nitrogen source (Nacry et al., 2013), indicating that *CYP735As* of rice and *Arabidopsis* respond to nitrogen in a way relevant to the nitrogen source availability of the plants. Nitrogen-dependent tZ-type CK accumulation has been proposed to be involved in growth enhancement in response to nitrogen supply in various plants (Takei et al., 2001b; Kamada-Nobusada et al., 2013; Ohashi et al., 2017; Maeda et al., 2018; Sakakibara, 2021; Shibasaki et al., 2021). Thus, the dual nitrogen-signal-dependent upregulation of tZ-type CK biosynthesis might play a similar role in rice. Consistent to that, there are reports suggesting that rice grows better with simultaneous supply of nitrate and ammonium than exclusive supply of nitrate or ammonium (Chanch and Ohira, 1982; Ying-Hua et al., 2006). Together, these differences between rice and *Arabidopsis* might be the result of evolution and/or domestication to optimize growth depending on their body plan and growth environments, though further investigation into the physiological function of tZ-type CKs in other plant species is necessary to clarify this issue.

## Materials and Methods

### Plant material and growth conditions

*Arabidopsis thaliana* ecotype Columbia (Col-0) was used as the wild type. The *cyp735a1-2 cyp735a2-2* double mutant was characterized previously (Kiba et al., 2013). *Arabidopsis* plants were grown on half-strength Murashige-Skoog (1/2x MS) agar plates (1% agar and 1% sucrose) placed vertically or on soil (Supermix A, Sakata, Japan) at 22°C under fluorescent light (100 μmol m^-2^ s^-1^, 16 h light/8 h dark).

*Oryza sativa* cultivar Nipponbare (NB) was used as the wild type. Rice seeds were germinated on moist filter paper at 30°C for 2-3 days in the dark and then grown under hydroponic culture or soil culture conditions. For hydroponic culture, germinated seeds were sown on mesh trays floating on the nutrient solution (Makino et al., 1985) with or without nitrogen source (NH_4_NO_3_ was replaced with KCl) and grown in an environment-controlled greenhouse at 28°C with a 15-h light (500 μmol m^-2^ s^-1^)/9-h dark cycle. The nutrient solution was renewed every 3-4 days. To analyze *CYP735A3* and *CYP735A4* expression in response to nitrogen, 12-day-old seedlings grown on the nutrient solution without nitrogen source were incubated with the nutrient solution without nitrogen source supplemented with KCl, NH_4_Cl, KNO_3_, or methionine sulfoximine (MSX). For soil culture, germinated seeds were sown on a synthetic soil (SunAgro, No. 3) and grown in an environment-controlled greenhouse at 28°C with a 15-h light (500 μmol m^-2^ s^-1^)/9-h dark cycle. Fifteen-day-old seedlings were transplanted in perforated plastic pots (10 × 12 × 15 cm) filled with nutrient-free soil (Aichi Medel) supplemented with slow-release fertilizer (N, P, and K at 0.3, 0.3, and 0.3 g/kg soil, respectively) and grown in a greenhouse at Nagoya University with supplemental lighting (13-h light/11-h dark cycle).

### Generation of transgenic lines overexpressing *CYP735A3* or *CYP735A4* in *cyp735a1 cyp735a2*

The *CYP735A3* and *CYP735A4* cDNA were amplified with the specific primer sets oxCYP735A3-F/R and oxCYP735A4-F/R, respectively (Table S4). The amplified fragments were cloned into the *Xba* I/*Sac* I site of pBI121 (Clontech) located downstream of the cauliflower mosaic virus (CaMV) 35S promoter and introduced into *cyp735a1-2 cyp735a2-2* double mutant by floral-dipping (Clough and Bent, 1998).

### CRISPR-Cas9 mutagenesis of *CYP735A3* and *CYP735A4*

The mutants of *CYP735A3* and *CYP735A4* were generated using the transfer RNA-based-multiplex CRISPR-Cas9 vector, pMgPoef4_129-2A-GFP (Toda et al., 2019). Guide sequences for *CYP735A3* and *CYP735A4* were designed by CHOPCHOP (Labun et al., 2019) and two multiplex CRISPR-Cas9 vectors pMg129_A3A4-1 and pMg129_A3A4-2, each harboring three guide sequences for *CYP735A3* and *CYP735A4* sequences, were constructed as described (Toda et al., 2019). pMg129_A3A4-1 and pMg129_A3A4-2 contained guide sequences g3-1, g4-1-1, and g4-1-2 (Fig. S1), and g3-2, g4-1-2, and g4-1-2 (Fig. S2), respectively. Transgenic rice plants were generated by the *Agrobacterium tumefaciens*-mediated method (Hirose et al., 2005) using EHA105 strain harboring the vectors. Mutations were identified by DNA sequencing of PCR products amplified with specific primer sets, gCYP735A3-F2/R2 and gCYP735A4-F3R3 (Table S4), and genomic DNA prepared from T1 plants. A T1 line transformed using pMg129_A3A4-2 but without any mutations in *CYP735A3* and *CYP735A* was used as the vector control line. For assays, *cyp735a3-1, cyp735a4-2, cyp735a3-1 cyp735a4-1, cyp735a3-2 cyp735a4-2* and vector control identified by genotyping progenies of these lines were used.

### Quantification of plant hormones

Cytokinin levels were determined as described previously (Kojima et al., 2009; Kojima and Sakakibara, 2012) using an ultra-performance liquid chromatograph coupled with a tandem quadrupole mass spectrometer (ACQUITY UPLC™ System/XEVO-TQS; Waters, Milford, MA, USA) with an octadecylsilyl (ODS) column (ACQUITY UPLC HSS T3, 1.8 μm, 2.1 mm × 100 mm, Waters). In the results, iP, iP-riboside, iP-riboside 5’-phosphates, iP-7-*N*-glucoside, and iP-9-*N*-glucoside are collectively referred to as “iP-type cytokinin”; tZ, tZ-riboside, tZ-riboside 5’-phosphates, tZ-7-*N*-glucoside, tZ-9-*N*-glucoside, tZ-*O*-glucoside, and tZR-*O*-glucoside are collectively referred to as “tZ-type cytokinin”; cZ, cZ-riboside, cZ-riboside 5’-phosphates, cZ-*O*-glucoside, and cZR-*O*-glucoside are collectively referred to as “cZ-type cytokinin”.

### Gene expression analysis

Total RNA was extracted from root and shoot samples using the RNeasy Plant Mini kit (QIAGEN) in combination with the RNase-Free DNase set (QIAGEN). Total RNA was used for first strand cDNA synthesis by the ReverTra Ace qPCR RT Master Mix (Toyobo). Quantitative reverse transcription-PCR (qRT-PCR) was performed on a Quant Studio 3 Real-Time PCR system (Thermo Fisher) with the KAPA SYBR Fast qPCR kit (KAPA Biosystems). *ZIURP1* (LOC_Os03g08010/Os03g0234200) was used as an internal control because this gene has been shown to be one of the most stably expressed genes in rice (Pabuayon et al., 2016). Primer sets are listed in Table S4.

### Promoter-GUS analysis

The *CYP735A3* promoter (proCYP735A3; -3636 to +21 bp relative to the inferred initiation codon) and *CYP735A4* promoter (proCYP735A4; -3623 to +21 bp) were amplified with PrimeSTAR GXL DNA polymerase (Takara) and the specific primer sets proCYP735A3-F/R and proCYP735A4-F/R, respectively (Table S4). The fragments were cloned into pENTR/D-TOPO vector (Thermo Fisher), sequenced, and then integrated into the GATEWAY binary vector pCAMBIA-GW-GUS (Kudo et al., 2012) to generate pCAMBIA1390-proCYP735A3:GUS and pCAMBIA1390-proCYP735A4:GUS vectors. Transgenic rice plants were generated by the *Agrobacterium tumefaciens*-mediated method (Hirose et al., 2005) using EHA105 strain harbouring the vectors. Histochemical analysis of GUS activity was conducted as described previously (Hirose et al., 2005; Kiba et al., 2018). Fresh and GUS-stained samples were fixed and then embedded in Technovit 7100 resin (Heraeus Kulzer) as instructed by the manufacturer. Sections were produced using a Leica RM2165 microtome (Leica) and observed under an Olympus BX51 microscope (Olympus).

### Morphological analyses

The rosette diameter, shoot length, root length, leaf blade length, leaf sheath length, leaf width, panicle length, and culm length were measured manually or determined from pictures using ImageJ (https://imagej.nih.gov/ij/). The days leading to the appearance of a panicle in main stem is defined as “days to heading”.

### Treatment with phytohormones and nitrogen sources

For phytohormone treatments, 12-day-old Nipponbare seedlings grown in hydroponic culture with the nutrient solution (Makino et al., 1985) containing 1 mM NH_4_NO_3_ were used. Phytohormone stock solutions were prepared by dissolving *N*^6^-(Δ^2^-isopentenyl)adenine (iP), *trans*-zeatin (tZ), indole-3-acetic acid (IAA), and abscisic acid (ABA) in dimethyl sulfoxide (DMSO) at 100 mM and added to the nutrient solution. Roots of the seedlings were submerged in the nutrient solution containing 1 μM iP, 1 μM tZ, 10 μM IAA, 50 μM ABA, 0.001% DMSO (mock for iP and tZ treatment), or 0.05% DMSO (mock for IAA and ABA treatment) and harvested after 2 h.

For nitrogen treatments, Nipponbare seedlings were grown in hydroponic culture with the nitrogen-free nutrient solution (1 mM NH_4_NO_3_ was replaced with 1 mM KCl) for 12 days. Roots of the seedlings were incubated with the nitrogen-free nutrient solution supplemented with 5 mM KCl, 5 mM NH_4_Cl, or 5 mM KNO_3_ for 2 h or 4 h. In case of MSX and nitrogen co-treatment, roots were pre-treated with the nitrogen-free nutrient solution containing 2 mM MSX for 2 h and then incubated with the MSX containing nitrogen-free nutrient solution supplemented with 5 mM KCl, 5 mM KNO_3_ or 50 mM glutamine for 2 h.

### Statistical analysis

Data are shown as means ± standard deviation (SD) of one representative experiment. In order to examine whether hormone concentration, gene expression, or morphological data were significantly different between treatments, Student’s *t*-test, and Tukey’s honestly significant difference (HSD) test were performed using KaleidaGraph ver. 4.1 software (Synergy Software).

### Accession numbers

Sequence data for the genes described in this article can be found in The Arabidopsis Information Resource database (http://www.arabidopsis.org) and Oryzabase (https://shigen.nig.ac.jp/rice/oryzabase/) under the following accession numbers: *CYP735A1* (At5g38450), *CYP735A2* (At1g67110), *ACT8* (AT1G49240), *CYP735A3* (LOC_Os08g33300/Os08g0429800), *CYP735A4* (LOC_Os09g23820/Os09g0403300), *ZIURP1* (LOC_Os03g08010/Os03g0234200), *OsRR1* (LOC_Os04g36070/Os04g0442300), *OsRR2* (LOC_Os02g35180/Os02g0557800), *OsRR4* (LOC_Os01g72330/Os01g0952500), *OsRR6* (LOC_Os04g57720/Os04g0673300), *OsRR9* (LOC_Os11g04720/Os11g0143300), *OsRR10* (LOC_Os12g04500/Os12g0139400), *OsRR21* (LOC_Os03g12350/Os03g0224200), *OsRR23* (LOC_Os02g55320/Os02g0796500), *OsSIPP2C1* (LOC_Os09g15670/Os09g0325700), *OsGH3*.*2* (LOC_Os01g55940/Os01g0764800), *OsIPT4* (LOC_Os03g59570/Os03g0810100), *OsNIA1* (LOC_Os08g36480/Os08g0468100).

## Acknowledgments

This work was supported by JSPS KAKENHI (No. JP18H04793 and JP20H02888 for T.K., JP17H06473 for H.S.).

## Author contributions

T.K. and H.S. designed the experiments; T.K., K.M., A.M., Y.T., M.K., T.H., Y.O., K.O., H.S. performed the experiments, T.K., K.M., A.M. analyzed the data, and T.K. and H.S. wrote the paper.

## Legends to Figures

**Supplemental Figure 1. Phylogenetic tree of *Arabidopsis* cytochrome P450s, CYP735A3 and CYP735A4**

(A) A phylogenetic tree of 293 *Arabidopsis* P450s, CYP735A3 (LOC_Os08g33300/Os08g0429800) and CYP735A4 (LOC_Os09g23820/Os09g0403300). (B) Enlargement of a subclade with CYP735A3 and CYP735A4. Full-length amino acid sequences were obtained from Plant P450 database (https://erda.dk/public/vgrid/PlantP450/index.html). A phylogenetic tree was inferred by the Neighbour-Joining method using MEGAX (https://www.megasoftware.net/). The tree is drawn to scale, with branch lengths in the same units as those of the evolutionary distances used to infer the phylogenetic tree. The evolutionary distances were computed using the JTT matrix-based method and are in the units of the number of amino acid substitutions per site. All positions containing gaps and missing data were eliminated. The values at the nodes indicate the bootstrap values (using 500 replications).

**Supplemental Figure 2. The *cyp735a3-1 cyp735a4-1* mutant generated by the CRISPR/Cas9 system**

(A) Schematic representation of CRISPR target sites to generate *cyp735a3-1* (*a3-1*) and *cyp735a4-1* (*a4-1*) alleles. Boxes represent exons; horizontal bars, introns; triangles, CRISPR target sites. The blue box represents the “A helix”. The bar indicates a 200 bp scale. (B) Partial sequences of wild-type *CYP735A3* and *cyp735a3-1* (*a3-1*) mutant. The sequence corresponding to a guide RNA used to generate the *a3-1* mutation (g3-1) is highlighted in yellow. (C) Partial sequences of wild-type *CYP735A4* and *cyp735a4-1* (*a4-1*) mutant. The sequence corresponding to guide RNAs used to generate the *a4-1* mutation (g4-1-1 and g4-1-2) is highlighted in yellow. Numbers in (B) and (C) represent positions in a genome sequence when the first nucleotide of the putative start codon is counted as 1. The red letter and red dash indicate an inserted and deleted sequence, respectively.

**Supplemental Figure 3. The *cyp735a3-2 cyp735a4-2* mutant generated by CRISPR/Cas9 system**

(A) Schematic representation of CRISPR target sites to generate *cyp735a3-2* (*a3-2*) and *cyp735a4-2* (*a4-2*) alleles. Boxes represent exons; horizontal bars, introns; triangles, CRISPR target sites. The blue box represents the “A helix”. The bar indicates a 200 bp scale. (B) Partial sequences of wild-type *CYP735A3* and *cyp735a3-2* (*a3-2*) mutant. The sequence corresponding to a guide RNA used to generate the *a3-2* mutation (g3-2) is highlighted in yellow. (C) Partial sequences of wild-type *CYP735A4* and *cyp735a4-2* (*a4-2*) mutant. The sequence corresponding to guide RNAs used to generate the *a4-2* mutation (g4-2-1 and g4-2-2) is highlighted in green. Numbers in (B) and (C) represent positions in a genome sequence when the first nucleotide of the putative start codon is counted as 1. The red letter and red dash indicate an inserted and deleted sequence, respectively.

**Supplemental Figure 4. Deduced amino acid sequences of *CYP735A3, cyp735a3-1* and *cyp735a3-2***

Deduced amino acid sequences of *CYP735A3, cyp735a3-1* (*a3-1*), and *cyp735a3-2* (*a3-2*) were aligned by Clustal Omega (https://www.ebi.ac.uk/Tools/msa/clustalo/). Triangles indicate CRISPR target sites.

**Supplemental Figure 5. Deduced amino acid sequences of *CYP735A4, cyp735a4-1* and *cyp735a4-2***

Deduced amino acid sequences of *CYP735A4, cyp735a4-1* (*a4-1*), and *cyp735a4-2* (*a4-2*) were aligned by Clustal Omega (https://www.ebi.ac.uk/Tools/msa/clustalo/). Triangles indicate CRISPR target sites. The “A helix” is highlighted in blue.

**Supplemental Figure 6. Effect of disruption of *CYP735A*s on vegetative growth**

(A) A representative image of Nipponbare (NB), vector control (VC), *cyp735a3-1 cyp735a4-1* (*a3a4-1*), and *cyp735a3-1* (*a3-1*) seedlings grown 14 days in hydroponic culture. Scale bar, 5 cm. (B) Quantification of shoot length, shoot fresh weight, root length, and root fresh weight of 14-day-old seedlings. NB, VC, *cyp735a3-1* (*a3-1*), and *a3a4-1* seedlings were grown in hydroponic culture. (C) A representative image of NB, VC, *a3-1, cyp735a4-1* (*a4-2*), *cyp735a3-2 cyp735a4-2* (*a3a4-2*) seedlings grown for 43 days on soil. Scale bar, 20 cm. (D) Plant height and leaf number of NB, VC, *a3-1, a4-2, a3a4-1*, and *cyp735a3-2 cyp735a4-2* (*a3a4-2*) grown for 30 days on soil. Error bars represent standard deviation of biological replicates (B, n=6-13; D, n=9-12). Different lowercase letters indicate statistically significant differences as indicated by Tukey’s HSD test (*p*<0.05).

