## Supplemental Figures for "The *trans*-zeatin-type side-chain modification of cytokinins controls rice growth"

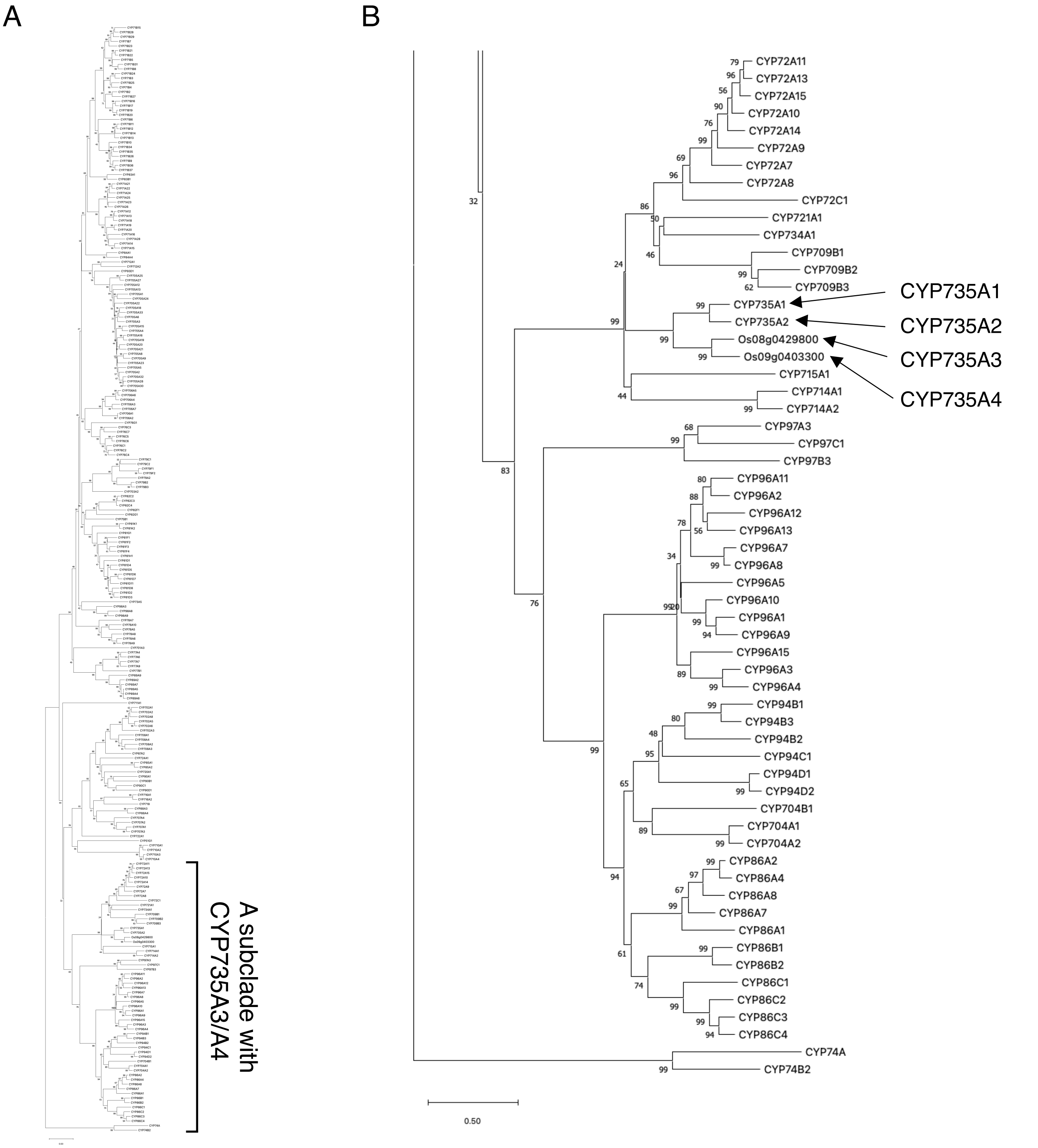


**Supplemental Figure 1. Phylogenetic tree of *Arabidopsis* cytochrome P450s, CYP735A3 and CYP735A4**
(A) A phylogenetic tree of 293 *Arabidopsis* P450s, CYP735A3 (LOC_Os08g33300/Os08g0429800) and CYP735A4 (LOC_Os09g23820/Os09g0403300). (B) Enlargement of a subclade with CYP735A3 and CYP735A4. Full-length amino acid sequences were obtained from Plant P450 database (https://erda.dk/public/vgrid/PlantP450/index.html). A phylogenetic tree was inferred by the Neighbour-Joining method using MEGAX (https://www.megasoftware.net/). The tree is drawn to scale, with branch lengths in the same units as those of the evolutionary distances used to infer the phylogenetic tree. The evolutionary distances were computed using the JTT matrix-based method and are in the units of the number of amino acid substitutions per site. All positions containing gaps and missing data were eliminated. The values at the nodes indicate the bootstrap values (using 500 replications).

A

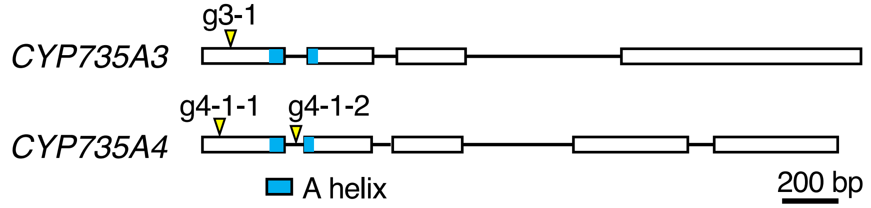


B

```
                          g3-1
CYP735A3:  79 ACCGTCTCATGCTACTACCTCA-CGCCGGCGAGG 111
a3-1     :    ACCGTCTCATGCTACTACCTCACCGCCGGCGAGG
```

C

```
                    g4-1-1
CYP735A4: 46   CTCGTGGCGCTGCTTCTGAGGGCGGCGTGGGTGACCCTGT 85
a4-1     :     CTCGTGGCGCTGCTTCTGA---------------------

CYP735A4: 86   CCTGCTACTGGCTGACGCCAATGAGGATCCGCCGCGCCAT 125
a4-1     :     ----------------------------------------

CYP735A4: 126 GGCGGCGCAGGGCGTGCGCGGCCCGCCGCCGCGCCCGCTC 165
a4-1     :     ----------------------------------------

CYP735A4: 166 GTCGGCAACCTCCGGGAGGTGTCGGCGCTCGTGGCGAGGG 205
a4-1     :     ----------------------------------------

CYP735A4: 206 CCACCGCCGACGACATGCCGTCCCTCAGCCACGACATCGT 245
a4-1     :     ----------------------------------------

CYP735A4: 246 CGGCCGCCTCATGCCCCACTACGTGCTCTGGTCCGGGACA 285
a4-1     :     ----------------------------------------

CYP735A4: 286 TACGGTGAGCACGCCACGCGCGACACGCCGCCGCCGGC 325
a4-1     :     ----------------------------------------
                  g4-1-2
CYP735A4: 326 GATTGTTCGTTCGTTCGTTGACTCTTTTTGGTTTGCAGGC 365
a4-1     :     --TTGTTCGTTCGTTCGTTGACTCTTTTTGGTTTGCAGGC
```

**Supplemental Figure 2. The *cyp735a3-1 cyp735a4-1* mutant generated by the CRISPR/Cas9 system**
(A) Schematic representation of CRISPR target sites to generate *cyp735a3-1* (*a3-1*) and *cyp735a4-1* (*a4-1*) alleles. Boxes represent exons; horizontal bars, introns; triangles, CRISPR target sites. The blue box represents the “A helix” . The bar indicates a 200 bp scale. (B) Partial sequences of wild-type *CYP735A3* and *cyp735a3-1* (*a3-1*) mutant. The sequence corresponding to a guide RNA used to generate the *a3-1* mutation (g3-1) is highlighted in yellow. (C) Partial sequences of wild-type *CYP735A4* and *cyp735a4-1* (*a4-1*) mutant. The sequence corresponding to guide RNAs used to generate the *a4-1* mutation (g4-1-1 and g4-1-2) is highlighted in yellow. Numbers in (B) and (C) represent positions in a genome sequence when the first nucleotide of the putative start codon is counted as 1. The red letter and red dash indicate an inserted and deleted sequence, respectively.

A

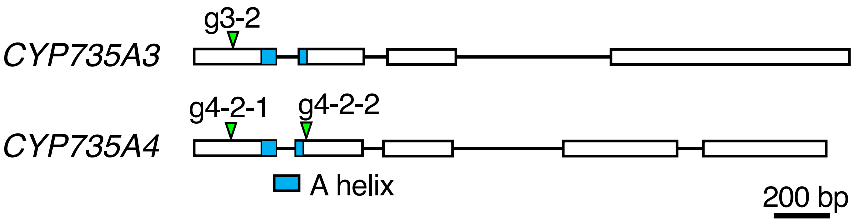


B

```
                                   g3-2
CYP735A3: 110 GGATCCGGCGGGTCCTGGCGAG-CCAGGGCGTGC 142
a3-2    :     GGATCCGGCGGGTCCTGGCGAGACCAGGGCGTGC
```

C

```
                           g4-2-1
CYP735A4: 86  CCTGCTACTGGCTGACGCCAATGAGGATCCGCCGCGCCAT 125
a4-2    :     CTCGTGGCGCTGCTTCTGA---------------------

CYP735A4: 126 GGCGGCGCAGGGCGTGCGCGGCCCGCCGCCGCGCCCGCTC 165
a4-2    :     ----------------------------------------

CYP735A4: 166 GTCGGCAACCTCCGGGAGGTGTCGGCGCTCGTGGCGAGGG 205
a4-2    :     ----------------------------------------

CYP735A4: 206 CCACCGCCGACGACATGCCGTCCCTCAGCCACGACATCGT 245
a4-2    :     ----------------------------------------

CYP735A4: 246 CGGCCGCCTCATGCCCCACTACGTGCTCTGGTCCGGGACA 285
a4-2    :     ----------------------------------------

CYP735A4: 286 TACGGTGAGCACGCCACGCGCGCGACACGCCGCCGCCGGC 325
a4-2    :     ----------------------------------------

CYP735A4: 326 GATTGTTCGTTCGTTCGTTGACTCTTTTTGGTTTGCAGGC 365
a4-2    :     ----------------------------------------

CYP735A4: 366 AAGCTGTTCGTGTACCTGTACGGGAGCGAGCCGAGGCTGT 405
a4-2    :     ------------------------------------CTGT
                  g4-2-2
CYP735A4: 406 GCCTGACTGACACCGCGCTGATCAAGGAGTTCTTGTCGTC 445
a4-2    :     GCCTGACTGACACCGCGCTCTGTGCCTGACTGACACCGCG
```

**Supplemental Figure 3. The *cyp735a3-2 cyp735a4-2* mutant generated by CRISPR/Cas9 system**
(A) Schematic representation of CRISPR target sites to generate *cyp735a3-2* (*a3-2*) and *cyp735a4-2* (*a4-2*) alleles. Boxes represent exons; horizontal bars, introns; triangles, CRISPR target sites. The blue box represents the “A helix”. The bar indicates a 200 bp scale. (B) Partial sequences of wild-type *CYP735A3* and *cyp735a3-2* (*a3-2*) mutant. The sequence corresponding to a guide RNA used to generate the *a3-2* mutation (g3-2) is highlighted in yellow. (C) Partial sequences of wild-type *CYP735A4* and *cyp735a4-2* (*a4-2*) mutant. The sequence corresponding to guide RNAs used to generate the *a4-2* mutation (g4-2-1 and g4-2-2) is highlighted in green. Numbers in (B) and (C) represent positions in a genome sequence when the first nucleotide of the putative start codon is counted as 1. The red letter and red dash indicate an inserted and deleted sequence, respectively.

```
                                                 g3-1      g3-2
                                                  ▼          ▼
CYP735A3        MAAAVLVAIALPVSLALLLVAKAVWVTVSCYYLTPARIRRVLASQG--------------      46
a3-1            MAAAVLVAIALPVSLALLLVAKAVWVTVSCYYLTAGEDPAGPGEPGRARPAAAAARRQPP      60
a3-2            MAAAVLVAIALPVSLALLLVAKAVWVTVSCYYLTPARIRRVLARPGRARPAAAAARRQPP      60
                ********************************** ..      . *

CYP735A3        --------------------VRGPPP---RPLVGNLRDVSALVAESTAADMASL-SH---      79
a3-1            RRVGARRRVHRRRHGLPQPRHRRPPPPPLRPLVQHVREAVRVLVERERAAGVRDGGRHGAG      120
a3-2            RRVGARRRVHRRRHGLPQPRHRRPPPPPLRPLVQHVREAVRVLVERERAAGVRDGGRHGAG      120
                                     * ***   **** ::*:.  ::... **.: .   *

CYP735A3        -DIVARLLPHYVLWSNTYGRRFVYWYGSEPRVCVTEAGMVRELLSSRHAHVTGKSWLQRQ      138
a3-1            APVVAARARHRQVVAAAAG------------------------------RQALHRPWPPHG      151
a3-2            APVVAARARHRQVVAAAAG------------------------------RQALHRPWPPHG      151
                  :**    *  : : : *                              :.  : *  :

CYP735A3        GAKHFIGRGLLMANGATWSHQRHVVAPAFMAD-RLKG---------RVGHMVECTRQTVRA      189
a3-1            -------------------QRRHLVAPAPRRRAGVHGRPAQGEGGAHGGVHEADGAGA--      190
a3-2            -------------------QRRHLVAPAPRRRAGVHGRPAQGEGGAHGGVHEADGAGA--      190
                                   ::**:****      ::*          * : *.    .

CYP735A3        LRDAVARSGNEVEIGAHMARLAGDVIARTEFDTSYETGKRIFLLIEE-LQRLTARSSRYL      248
a3-1            -----EG------CGGEVRERGGDRRAHGEARRRRDRAHRVRHELRDRQEDLPPHR--GA      237
a3-2            -----EG------CGGEVRERGGDRRAHGEARRRRDRAHRVRHELRDRQEDLPPHR--GA      237
                              *..: . .**  *: *     : .:*:   :.:  : *  :

CYP735A3        WVPGSQYFPSKYRREIKRLNGELERLLKESIDRSRE--------IADEG---RTPSASPCG      298
a3-1            PAPHRPLQPLP--------LGPRQPVFSEQVQERDKAAERRAGAAAQGVHRPEPGDRRRG      289
a3-2            PAPHRPLQPLP--------LGPRQPVFSEQVQERDKAAERRAGAAAQGVHRPEPGDRRRG      289
                 .*     *           *  : ::.*.::.  :        * :*     *.    *

CYP735A3        RGLLGMLLAEMEKKEAGGNGGGE---VGYDAQMMIDECKTFFFAGHETSALLLTWAIMLL      355
a3-1            PDAVGVAVRPWPPRHAAGRDGEEGGRRQWRRRGRVRRPD----DDRRVQDLLLRRPR-DV      344
a3-2            PDAVGVAVRPWPPRHAAGRDGEEGGRRQWRRRGRVRRPD----DDRRVQDLLLRRPR-DV      344
                 . :*: :     :.*.*..* *     :  :  : . .     .:... ***      :

CYP735A3        ATHPAWQDKARAEVAAVCGGGAPSPDSLPKLAVLQMVINETLRLYPPAT-LLPRMAFEDI      414
a3-1            GAAPHLGHHAARHAPGVAGQGARRGRR--RLRRR-----RAVAGQPPEARRAPDGDQ---      394
a3-2            GAAPHLGHHAARHAPGVAGQGARRGRR--RLRRR-----RAVAGQPPEARRAPDGDQ---      394
                .: *   .:*  .. .*.* **       :*        .::   ** :   *

CYP735A3        ELGGGALRVPSGASVWIPVLAIH------HDEGAWGRD-----AHEFRPDRFAPGRPRPP      463
a3-1            ---RDAAAVPAGDAAAADGVRGHRARRGRAPGAEWRVGVDPGARHPPRRGRVGPRRARVQ      451
a3-2            ---RDAAAVPAGDAAAADGVRGHRARRGRAPGAEWRVGVDPGARHPPRRGRVGPRRARVQ      451
                    .*  **:* :.    :  *         . *  .      *  * .*..* * *

CYP735A3        AGAFLPFAAGPRNCVGQAYAMVEAKVALAMLLSSFRF-AISDEYRHAPVNVLTLRPRHGV      522
a3-1            AGQVR---AGTAAAAGGGVPAVRRRAAQ-LRRAGVRHGGGQGRARHAPLQL-PLRHLRRV      506
a3-2            AGQVR---AGTAAAAGGGVPAVRRRAAQ-LRRAGVRHGGGQGRARHAPLQL-PLRHLRRV      506
                ** .    **   ..* .   *. :.*  :  :..*. . ... ****:::  **  : *

CYP735A3        PVRL-----LP------LPPPRP-------   534
a3-1            PARAGERAHAPATPRRARPPPAAAAAAPIX   536
a3-2            PARAGERAHAPATPRRARPPPAAAAAAPIX   536
                *.*       *       ***
```

**Supplemental Figure 4. Deduced amino acid sequences of *CYP735A3*, *cyp735a3-1* and *cyp735a3-2***
Deduced amino acid sequences of *CYP735A3*, *cyp735a3-1* (*a3-1*), and *cyp735a3-2* (*a3-2*) were aligned by Clustal Omega (https://www.ebi.ac.uk/Tools/msa/clustalo/). Triangles indicate CRISPR target sites.

```
                                         g4-1-1          g4-2-1
                                            ▼               ▼
CYP735A4       MAVLVSLMVIAASSPLVALLLRAAWVTLSCYWLTPMRIRRAMAAQGVRGPPPRPLVGNLR     60
a4-1           MAVLVSLMVIAASSPLVALLLIVRSF-VDS------------------------------     17
a4-2           MAVLVSLMVIAASSPLVALLLRAAWVTLSCYWLTPVPD----------------------     38
               *****************

                                                 g4-1-2            g4-2-2
                                                    ▼                  ▼
CYP735A4       EVSALVARATADDMPSLSHDIVGRLMPHYVLWSGTYGKLFVYLYGSEPRLCLTDTALIKE     120
a4-1           ------------------------------FW--FAGKLFVYLYGSEPRLCLTDTALIKE     45
a4-2           ------------------------------------------------------------     38


CYP735A4       FLSSKYAHATGKSWLQRQGTKHFIGGGLLMANGARWAHQRHVVAPAFMADKLKARGRVGR     180
a4-1           FLSSKYAHATGKSWLQRQGTKHFIGGGLLMANGARWAHQRHVVAPAFMADKLKARGRVGR     105
a4-2           ------------------------------------------------------------     38

CYP735A4       MVECTKQAIRELRDAAAGRRGEEVEIGAHMTRLTGDIISRTEFNTSYDTGKRIFLLLEHL     240
a4-1           MVECTKQAIRELRDAAAGRRGEEVEIGAHMTRLTGDIISRTEFNTSYDTGKRIFLLLEHL     165
a4-2           ------------------------------------------------------------     38

CYP735A4       QRLTSRSSRHLWIPGSQYFPSKYRREIRRLNGELEAVLMESIRRSREIADEGRAAVATYG     300
a4-1           QRLTSRSSRHLWIPGSQYFPSKYRREIRRLNGELEAVLMESIRRSREIADEGRAAVATYG     225
a4-2           ------------------------------------------------------------     38

CYP735A4       RGLLAMLLSEMEEKEKNGGGGGGEFSYDAQLVIDECKTFFFAGHETSALLLTWAIMLLAT     360
a4-1           RGLLAMLLSEMEEKEKNGGGGGGEFSYDAQLVIDECKTFFFAGHETSALLLTWAIMLLAT     285
a4-2           ------------------------------------------------------------     38

CYP735A4       NPAWQEKARTEVAAVCGDHPPSADHLSKLTVLQMIIQETLRLYPPATLLPRMAFEDIQLG     420
a4-1           NPAWQEKARTEVAAVCGDHPPSADHLSKLTVLQMIIQETLRLYPPATLLPRMAFEDIQLG     345
a4-2           ------------------------------------------------------------     38

CYP735A4       GLRLPRGLSVWIPVLAIHHDESIWGPDAHEFRPERFAPGARRPSAAGAARFLPFAAGPRN     480
a4-1           GLRLPRGLSVWIPVLAIHHDESIWGPDAHEFRPERFAPGARRPSAAGAARFLPFAAGPRN     405
a4-2           ------------------------------------------------------------     38

CYP735A4       CVGQAYALVEAKVVLAMLLSAFRFAISDNYRHAPENVLTLRPKHGVPVHLRPLRP     535
a4-1           CVGQAYALVEAKVVLAMLLSAFRFAISDNYRHAPENVLTLRPKHGVPVHLRPLRP     460
a4-2           -------------------------------------------------------     38
```

**Supplemental Figure 5. Deduced amino acid sequences of *CYP735A4*, *cyp735a4-1* and *cyp735a4-2***
Deduced amino acid sequences of *CYP735A4*, *cyp735a4-1* (*a4-1*), and *cyp735a4-2* (*a4-2*) were aligned by Clustal Omega (https://www.ebi.ac.uk/Tools/msa/clustalo/). Triangles indicate CRISPR target sites. The “A helix is highlighted in blue.

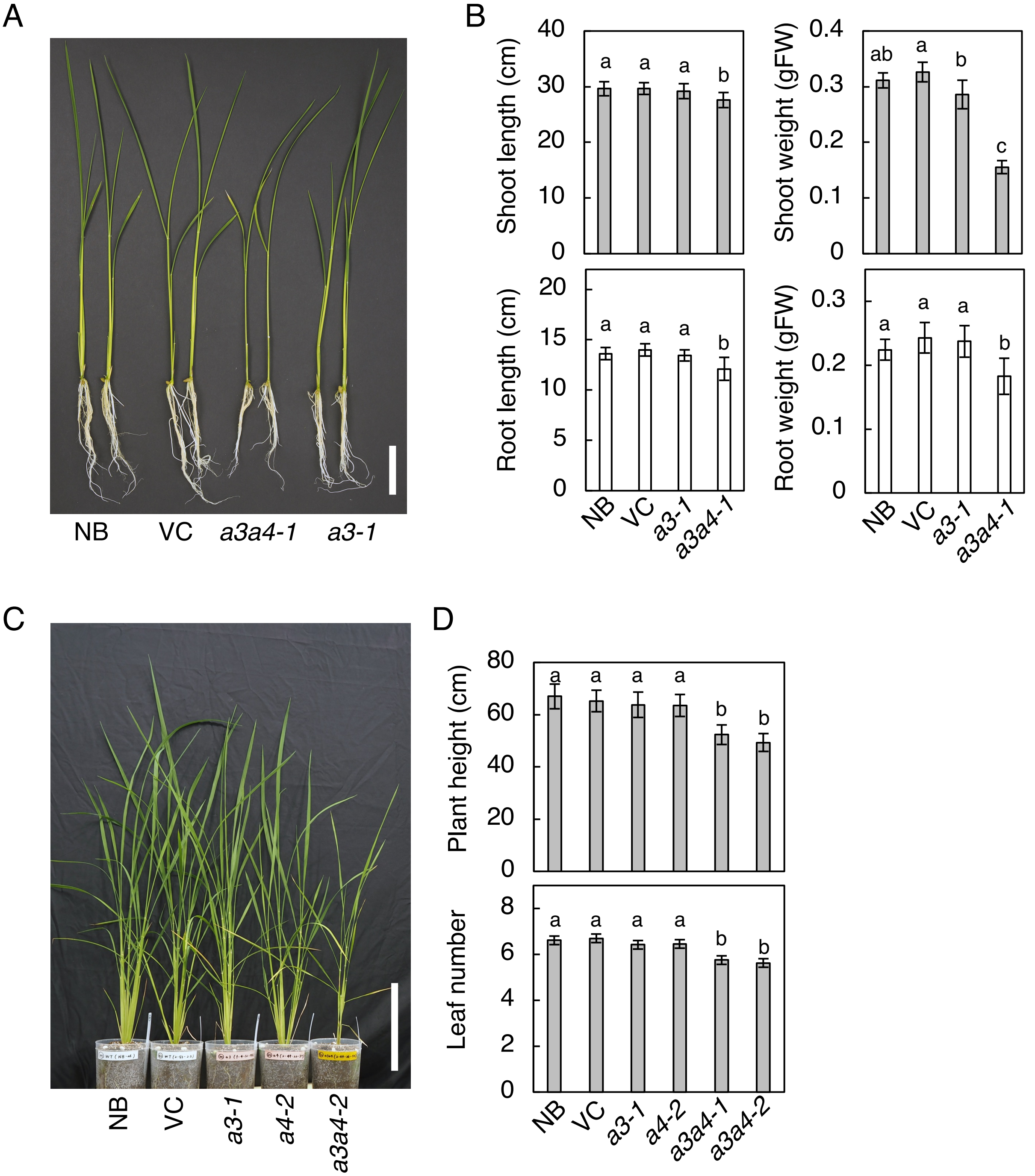


**Supplemental Figure 6. Effect of disruption of *CYP735A*s on vegetative growth**

(A) A representative image of Nipponbare (NB), vector control (VC), *cyp735a3-1 cyp735a4-1* (*a3a4-1*), and *cyp735a3-1* (*a3-1*) seedlings grown 14 days in hydroponic culture. Scale bar, 5 cm. (B) Quantification of shoot length, shoot fresh weight, root length, and root fresh weight of 14 day-old seedlings. NB, VC, *cyp735a3-1* (*a3-1*), and *a3a4-1* seedlings were grown in hydroponic culture. (C) A representative image of NB, VC, *a3-1*, *cyp735a4-1* (*a4-2*), *cyp735a3-2 cyp735a4-2* (*a3a4-2*) seedlings grown for 43 days on soil. Scale bar, 20 cm. (D) Plant height and leaf number of NB, VC, *a3-1*, *a4-2*, *a3a4-1*, and *cyp735a3-2 cyp735a4-2* (*a3a4-2*) grown for 30 days on soil. Error bars represent standard deviation of biological replicates (B, n=6-13; D, n=9-12). Different lowercase letters indicate statistically significant differences as indicated by Tukey's HSD test ($p$<0.05).
