## Supplemental Tables for "The *trans*-zeatin-type side-chain modification of cytokinins controls rice growth"

**Table S1. Cytokinin concentrations in Col-0, *cyp735a1 cyp735a2* mutant, and transgenic plants constitutively expressing *CYP735A3* (*A3*-ox) or *CYP735A4* (*A4*-ox) in *cyp735a1 cyp735a2* background.**

| pmol/gFW | Col-0 | *cyp735a1 cyp735a2* | *A3*-ox (Line 2) | *A4*-ox (Line 2) |
|---|---|---|---|---|
| tZ | 0.60 ± 0.10* | 0.05 ± 0.01 | 32.83 ± 13.72* | 48.64 ± 4.57* |
| tZR | 1.62 ± 0.23* | 0.07 ± 0.02 | 15.37 ± 3.73* | 21.59 ± 2.86* |
| tZRPs | 20.02 ± 2.11* | 1.01 ± 0.10 | 17.42 ± 3.20* | 19.77 ± 1.51* |
| cZ | 0.12 ± 0.01 | 0.2 ± 0.07 | 0.62 ± 0.49 | 0.78 ± 0.27 |
| cZR | 0.57 ± 0.15 | 0.61 ± 0.15 | 2.54 ± 2.50 | 4.10 ± 1.51 |
| cZRPs | 4.19 ± 0.5 | 4.72 ± 1.23 | 2.71 ± 1.33 | 2.42 ± 1.13 |
| iP | N.D. | 0.12 ± 0.00 | N.D. | N.D. |
| iPR | 0.66 ± 0.09 | 0.98 ± 0.26 | N.D. | N.D. |
| iPRPs | 33.48 ± 5.59* | 68.77 ± 12.63 | 2.97 ± 0.99* | 2.08 ± 0.11* |
| tZ7G | 18.95 ± 0.37* | 1.56 ± 0.11 | 31.5 ± 14.94* | 19.46 ± 1.13* |
| tZ9G | 8.93 ± 0.83* | 0.33 ± 0.07 | 16.81 ± 9.75* | 8.40 ± 0.57* |
| tZOG | 12.82 ± 1.22* | 0.75 ± 0.13 | 17.30 ± 8.24* | 9.54 ± 0.95* |
| cZOG | 2.8 ± 0.5* | 5.03 ± 1.27 | 0.95 ± 0.75* | 0.56 ± 0.30* |
| tZROG | 0.79 ± 0.12* | 0.05 ± 0.02 | 1.55 ± 0.87* | 0.94 ± 0.05* |
| cZROG | 1.09 ± 0.05 | 1.18 ± 0.20 | 0.75 ± 0.39 | 0.6 ± 0.32 |
| tZRPsOG | 0.38 ± 0.08 | N.D. | 0.53 ± 0.27 | 0.31 ± 0.08 |
| cZRPsOG | N.D. | N.D. | N.D. | N.D. |
| iP7G | 15.17 ± 0.6* | 35.66 ± 1.40 | 1.13 ± 0.51* | 0.73 ± 0.03* |
| iP9G | 2.78 ± 0.10* | 7.39 ± 0.22 | 0.15 ± 0.11* | 0.07 ± 0.01* |
| iP-type | 52.08 ± 5.89* | 112.92 ± 12.58 | 4.25 ± 1.61* | 2.89 ± 0.12* |
| tZ-type | 64.1 ± 3.41* | 3.81 ± 0.32 | 133.3 ± 14.42* | 128.67 ± 9.51* |
| cZ-type | 8.76 ± 0.90 | 11.73 ± 2.71 | 7.42 ± 2.71 | 8.47 ± 2.41 |
| Total CK | 124.94 ± 7.37 | 128.45 ± 12.5 | 144.97 ± 13.69 | 140.03 ± 7.17 |

Seedlings were grown for 12 days on 1/2x MS agart plates and whole seedlings were harvested. Data are means ± standard deviation (n = 4). gFW, gram fresh weight; *cyp735a1 cyp735a2, cyp735a1-2 cyp735a2-2*; tZ, *trans*-zeatin; tZR ,tZ riboside; tZRPs, tZ ribotides; cZ, *cis*-zeatin; cZR, cZ riboside; cZRPs, cZ ribotides; iP, *N*6-(*Δ*2-isopentenyl)adenine; iPR, iP riboside; iPRPs, iP ribotides; tZ7G, tZ-7-*N*-glucoside; tZ9G, tZ-9-*N*-glucoside; tZOG, tZ-*O*-glucoside; cZOG, cZ-*O*-glucoside; tZROG, tZR-*O*-glucoside; cZROG, cZR-*O*-glucoside; iP7G, iP-7-*N*-glucoside; iP9G, iP-9-*N*-glucoside, N.D., under the quantification detection limit.. *, significantly different from *cyp735a1 cyp735a2* in Student's *t*-test ($p$<0.05)

**Table S2. Cytokinin concentrations in Nipponbare, vector control, *cyp735a3-1* single mutant, and *cyp735a3-1 cyp735a4-1* double mutant.**

| | Shoot | | | | Root | | | |
|---|---|---|---|---|---|---|---|---|
| pmol/gFW | Nipponbare | Vector control | *cyp735a3-1* | *cyp735a3-1 cyp735a4-1* | Nipponbare | Vector control | *cyp735a3-1* | *cyp735a3-1 cyp735a4-1* |
| tZ | 0.34 ± 0.15 | 0.34 ± 0.16 | 0.15 ± 0.17 | N.D. | 0.04 ± 0.05 | 0.14 ± 0.05 | 0.08 ± 0.06 | N.D. |
| tZR | 0.12 ± 0.07 | 0.09 ± 0.06 | 0.09 ± 0.07 | 0.01 ± 0.02* | 0.07 ± 0.02 | 0.04 ± 0.01 | 0.04 ± 0.01 | 0.01 ± 0.01* |
| tZRPs | 0.1 ± 0.08 | 0.12 ± 0.04 | 0.08 ± 0.04 | N.D. | 0.35 ± 0.14 | 0.33 ± 0.07 | 0.27 ± 0.11 | 0.03 ± 0.03* |
| cZ | 1.28 ± 0.14 | 2.43 ± 1.45 | 1.62 ± 0.49 | 2.15 ± 0.84 | 2.15 ± 0.67 | 1.67 ± 0.64 | 2.19 ± 0.88 | 2.04 ± 0.23 |
| cZR | 1.53 ± 0.28 | 2.81 ± 1.81 | 1.69 ± 0.35 | 1.87 ± 0.34 | 1.99 ± 0.56 | 1.62 ± 0.42 | 1.99 ± 0.66 | 2.08 ± 0.55 |
| cZRPs | 0.62 ± 0.14 | 0.86 ± 0.10 | 0.74 ± 0.16 | 1.12 ± 0.43 | 2.59 ± 0.59 | 2.53 ± 0.62 | 3.36 ± 1.34 | 2.85 ± 0.97 |
| iP | 0.40 ± 0.06 | 0.36 ± 0.06 | 0.39 ± 0.11 | 0.69 ± 0.27 | 0.28 ± 0.07 | 0.23 ± 0.05 | 0.2 ± 0.02 | 0.28 ± 0.05 |
| iPR | 0.06 ± 0.00 | 0.24 ± 0.25 | 0.08 ± 0.03 | 0.11 ± 0.04 | 0.30 ± 0.09 | 0.28 ± 0.04 | 0.42 ± 0.08 | 0.93 ± 0.31* |
| iPRPs | 0.56 ± 0.08 | 0.46 ± 0.1 | 0.49 ± 0.12 | 1.13 ± 0.31* | 1.47 ± 0.25 | 1.33 ± 0.2 | 2.02 ± 0.64 | 2.37 ± 0.33* |
| tZ7G | N.D. | N.D. | N.D. | N.D. | N.D. | 0.01 ± 0.02 | N.D. | N.D. |
| tZ9G | 7.11 ± 1.89 | 9.19 ± 2.69 | 5.57 ± 2.48 | 0.07 ± 0.02* | 5.07 ± 1.26 | 7.18 ± 2.26 | 4.29 ± 0.87 | 0.05 ± 0.01* |
| tZOG | 0.18 ± 0.04 | 0.13 ± 0.10 | 0.24 ± 0.03 | 0.32 ± 0.03* | 0.14 ± 0.03 | 0.13 ± 0.03 | 0.13 ± 0.05 | 0.13 ± 0.07 |
| cZOG | 318.42 ± 20.50 | 310.04 ± 101.95 | 326.98 ± 33.68 | 291.78 ± 42.49 | 154.34 ± 5.81 | 147.97 ± 19.26 | 165.49 ± 29.53 | 126.67 ± 24.20 |
| tZROG | 0.1 ± 0.02 | 0.09 ± 0.01 | 0.08 ± 0.03 | 0.11 ± 0.03 | 0.29 ± 0.07 | 0.27 ± 0.05 | 0.23 ± 0.04 | 0.19 ± 0.03 |
| cZROG | 40.34 ± 0.86 | 36.09 ± 2.53 | 38.38 ± 1.98 | 37.19 ± 2.74 | 64.04 ± 3.96 | 60.04 ± 6.87 | 66.87 ± 11.01 | 72.92 ± 7.60 |
| tZRPsOG | N.D. | N.D. | N.D. | N.D. | N.D. | N.D. | N.D. | N.D. |
| cZRPsOG | 2.71 ± 0.25 | 2.87 ± 0.25 | 2.52 ± 0.10 | 3.16 ± 0.36 | 2.37 ± 0.58 | 2.58 ± 0.75 | 2.94 ± 0.33 | 2.87 ± 0.46 |
| iP7G | N.D. | N.D. | N.D. | N.D. | N.D. | 0.05 ± 0.08 | N.D. | N.D. |
| iP9G | 0.94 ± 0.13 | 1.17 ± 0.13 | 1.25 ± 0.31 | 2.22 ± 0.58* | 1.69 ± 0.18 | 1.91 ± 0.46 | 1.76 ± 0.26 | 3.76 ± 1.01* |
| tZ-type | 7.96 ± 2.07 | 9.97 ± 3.00 | 6.20 ± 2.59 | 0.51 ± 0.07* | 5.96 ± 1.35 | 8.10 ± 2.36 | 5.04 ± 0.93 | 0.41 ± 0.11* |
| iP-type | 1.95 ± 0.19 | 2.22 ± 0.40 | 2.22 ± 0.52 | 4.14 ± 1.00* | 3.74 ± 0.52 | 3.80 ± 0.63 | 4.41 ± 0.87 | 7.34 ± 1.51* |
| cZ-type | 364.88 ± 19.89 | 355.1 ± 108.00 | 371.93 ± 34.80 | 337.27 ± 41.77 | 227.48 ± 9.92 | 216.39 ± 24.81 | 242.86 ± 40.34 | 209.42 ± 28.31 |
| iP-type + tZ-type | 9.91 ± 2.24 | 12.19 ± 3.30 | 8.42 ± 2.94 | 4.65 ± 1.04* | 9.7 ± 1.85 | 11.91 ± 2.63 | 9.45 ± 1.51 | 7.76 ± 1.55 |
| Total CK | 374.79 ± 18.04 | 367.29 ± 110.59 | 380.35 ± 36.77 | 341.92 ± 42.58 | 237.18 ± 11.31 | 228.29 ± 24.84 | 252.3 ± 41.73 | 217.18 ± 28.65 |

Seedlings were grown hydroponically for 15 days and shoots and roots were harvested separately. Data are means ± standard deviation (n = 3-6). gFW, gram fresh weight; , tZ, *trans*-zeatin; tZR ,tZ riboside; tZRPs, tZ ribotides; cZ, cis-zeatin; cZR, cZ riboside; cZRPs, cZ ribotides; DZ, dihydrozeation; iP, *N*6-(Δ2-isopentenyl)adenine; iPR, iP riboside; iPRPs, iP ribotides; tZ7G, tZ-7-*N*-glucoside; tZ9G, tZ-9-*N*-glucoside; tZOG, tZ-*O*-glucoside; cZOG, cZ-*O*-glucoside; tZROG, tZR-*O*-glucoside; cZROG, cZR-*O*-glucoside; DZ9G, DZ-9-*N*-glucoside; iP7G, iP-7-*N*-glucoside; iP9G, iP-9-*N*-glucoside, N.D., under the quantification detection limit. *, significantly different from Nipponbare in Student's *t*-test ($p<0.05$)

**Table S3. Cytokinin concentrations in Nipponbare, vector control, *cyp735a4-2* single mutant, and *cyp735a3-2 cyp735a4-2* double mutant.**

| | Shoot | | | | Root | | | |
|---|---|---|---|---|---|---|---|---|
| pmol/gFW | Nipponbare | Vector control | *cyp735a4-2* | *cyp735a3-2 cyp735a4-2* | Nipponbare | Vector control | *cyp735a4-2* | *cyp735a3-2 cyp735a4-2* |
| tZ | 0.37 ± 0.05 | 0.48 ± 0.11 | 0.34 ± 0.13 | N.D. | 0.12 ± 0.02 | 0.07 ± 0.06 | 0.04 ± 0.04** | N.D. |
| tZR | 0.18 ± 0.06 | 0.29 ± 0.09 | 0.15 ± 0.09 | 0.01 ± 0.02* | 0.05 ± 0.03 | 0.05 ± 0.05 | 0.02 ± 0.02 | N.D. |
| tZRPs | 0.88 ± 0.23 | 0.86 ± 0.18 | 0.85 ± 0.18 | 0.03 ± 0.04* | 0.38 ± 0.13 | 0.38 ± 0.23 | 0.20 ± 0.10 | 0.08 ± 0.06 |
| cZ | 5.67 ± 10.21 | 1.35 ± 0.31 | 1.55 ± 0.2 | 1.31 ± 0.28 | 0.45 ± 0.19 | 0.30 ± 0.06 | 0.56 ± 0.51 | 0.34 ± 0.15 |
| cZR | 3.2 ± 2.03 | 2.13 ± 0.21 | 2.41 ± 0.45 | 2.41 ± 0.23 | 1.61 ± 0.40 | 1.55 ± 0.15 | 1.99 ± 0.47 | 1.88 ± 0.19* |
| cZRPs | 1.22 ± 0.41 | 1.16 ± 0.22 | 1.22 ± 0.15 | 1.56 ± 0.18 | 3.62 ± 2.65 | 2.22 ± 0.42 | 3.41 ± 2.05 | 3.34 ± 0.92 |
| iP | 0.71 ± 0.4 | 1.27 ± 0.73 | 1.77 ± 0.74* | 1.25 ± 0.32* | N.D. | N.D. | N.D. | 0.04 ± 0.1 |
| iPR | 0.29 ± 0.13 | 0.57 ± 0.31 | 0.45 ± 0.14* | 0.66 ± 0.21* | 0.32 ± 0.08 | 0.17 ± 0.06* | 0.57 ± 0.22 | 0.98 ± 0.29* |
| iPRPs | 1.34 ± 0.33 | 1.75 ± 0.7 | 2.7 ± 0.71* | 3.65 ± 0.61* | 0.67 ± 0.13 | 0.61 ± 0.2 | 1.44 ± 0.43* | 2.02 ± 0.47* |
| tZ7G | N.D. | N.D. | N.D. | N.D. | N.D. | N.D. | N.D. | N.D. |
| tZ9G | 11.22 ± 1.23 | 9.68 ± 0.81* | 5.85 ± 0.95* | 0.11 ± 0.08* | 4.50 ± 0.67 | 4.11 ± 1.06 | 2.26 ± 0.96* | 0.08 ± 0.03* |
| tZOG | 0.17 ± 0.08 | 0.17 ± 0.07 | 0.22 ± 0.06 | 0.21 ± 0.2 | 0.03 ± 0.05 | 0.21 ± 0.06* | 0.14 ± 0.13 | 0.09 ± 0.08* |
| cZOG | 223.14 ± 52.14 | 228.01 ± 45.08 | 250.02 ± 46.71 | 270.67 ± 34.6 | 184.07 ± 12.17 | 166.69 ± 11.65* | 197.41 ± 33.84 | 144.06 ± 59.64 |
| tZROG | 0.03 ± 0.05 | 0.02 ± 0.04 | 0.07 ± 0.07 | 0.09 ± 0.06 | 0.25 ± 0.12 | 0.32 ± 0.04 | 0.24 ± 0.06 | 0.22 ± 0.04* |
| cZROG | 26.66 ± 2.94 | 26.46 ± 2.72 | 28.32 ± 3.84 | 31.67 ± 2.88* | 52.24 ± 3.32 | 54.92 ± 3.81 | 56.25 ± 4.71 | 58.68 ± 4.39 |
| tZRPsOG | N.D. | N.D. | N.D. | N.D. | N.D. | N.D. | N.D. | N.D. |
| cZRPsOG | 2.38 ± 0.23 | 3.06 ± 0.59* | 3.34 ± 0.45* | 3.77 ± 0.77* | 2.48 ± 0.33 | 3.82 ± 0.91 | 4.25 ± 1.24* | 4.94 ± 0.92* |
| iP7G | N.D. | N.D. | N.D. | N.D. | N.D. | N.D. | N.D. | N.D. |
| iP9G | 3.78 ± 0.51 | 4.09 ± 0.95 | 5.12 ± 0.8* | 6.21 ± 1.07* | 1.48 ± 0.24 | 1.23 ± 0.13 | 2.19 ± 0.58 | 4.44 ± 1.18* |
| tZ-type | 12.86 ± 1.48 | 11.5 ± 0.70 | 7.48 ± 1.04* | 0.45 ± 0.26* | 5.33 ± 0.90 | 5.15 ± 0.94 | 2.89 ± 1.09* | 0.47 ± 0.13* |
| iP-type | 6.11 ± 1.23 | 7.69 ± 2.29 | 10.04 ± 2.12* | 11.77 ± 1.12* | 2.48 ± 0.42 | 2.00 ± 0.28 | 4.20 ± 1.19* | 7.48 ± 1.72* |
| cZ-type | 262.26 ± 52.20 | 262.17 ± 47.75 | 286.87 ± 45.62 | 311.38 ± 32.49 | 244.48 ± 12.02 | 229.49 ± 12.73 | 263.87 ± 38.36 | 213.25 ± 57.99 |
| iP-type + tZ-type | 18.98 ± 1.85 | 19.18 ± 1.29 | 17.52 ± 2.00 | 12.22 ± 1.30* | 7.81 ± 1.29 | 7.15 ± 1.07 | 7.09 ± 2.21 | 7.95 ± 1.82 |
| Total | 281.24 ± 53.55 | 281.35 ± 49.12 | 304.39 ± 46.95 | 323.6 ± 31.66 | 252.29 ± 12.42 | 236.64 ± 12.00 | 270.97 ± 39.97 | 221.2 ± 56.69 |

Seedlings were grown hydroponically for 13 days and shoots and roots were harvested separately. Data are means ± standard deviation (n = 6-8). gFW, gram fresh weight; , tZ, *trans*-zeatin; tZR ,tZ riboside; tZRPs, tZ ribotides; cZ, *cis*-zeatin; cZR, cZ riboside; cZRPs, cZ ribotides; DZ, dihydrozeation; iP, *N*6-(Δ2-isopentenyl)adenine; iPR, iP riboside; iPRPs, iP ribotides; tZ7G, tZ-7-*N*-glucoside; tZ9G, tZ-9-*N*-glucoside; tZOG, tZ-*O*-glucoside; cZOG, cZ-*O*-glucoside; tZROG, tZR-*O*-glucoside; cZROG, cZR-*O*-glucoside; DZ9G, DZ-9-*N*-glucoside; iP7G, iP-7-*N*-glucoside; iP9G, iP-9-*N*-glucoside, N.D., under the quantification detection limit. *, significantly different from Nipponbare in Student's *t*-test ($p<0.05$)

**Table S4. List of primers used for vector construction and genotyping.**

| Name (Forward/Reverse) | Purpose | Forward (5' to 3') | Reverse (5' to 3') |
|---|---|---|---|
| oxCYP735A3-F/R | Construction of pBI121-CYP735A3 | TTTCTAGATGGCAATGGCCGCCGCCGT | CTATGGGCGCGGCGGCGGCAGCGG |
| oxCYP735A4-F/R | Construction of pBI121-CYP735A4 | TTTCTAGATGGCGGTCCTCGTGTCGCTCA | CTATGGCCGCAGCGGCCGGAGG |
| gCYP735A3A4-1-1-F/R | Construction of pMgPoef4_129-2A-GFP-cyp735a3a4-1 | TTGGGTCTCGTGCACTCATGCTACTACCTCACGCGTTTTAGAGCTAGAAATAGCA | TTGGGTCTCCAGCAGCGCCACGTGCACCAGCCGGGAATCGAA |
| gCYP735A3A4-1-2-F/R | Construction of pMgPoef4_129-2A-GFP-cyp735a3a4-1 | TTGGGTCTCGTGCTTCTGAGGGGTTTTAGAGCTAGAAATAGCA | TTGGGTCTCCAAACGCGATTGTTCGTTCGTTCGTTGCACCAGCCGGGAATCGAA |
| gCYP735A3A4-2-1-F/R | Construction of pMgPoef4_129-2A-GFP-cyp735a3a4-2 | TTGGGTCTCGTGCACGGCGGGTCCTGGCGAGCCAGTTTTAGAGCTAGAAATAGCA | TTGGGTCTCCTCAGCCAGTAGCTGCACCAGCCGGGAATCGAA |
| gCYP735A3A4-2-2-F/R | Construction of pMgPoef4_129-2A-GFP-cyp735a3a4-2 | TTGGGTCTCGCTGACGCCAATGGTTTTAGAGCTAGAAATAGCA | TTGGGTCTCCAAACAGGCTGTGCCTGACTGACACTGCACCAGCCGGGAATCGAA |
| gCYP735A3-F2/R2 | Genotyping of *cyp735a3* mutations | TCCTCGTCGCCATCGCATTG | TTGCTCGTGAATGGGAGGAGC |
| gCYP735A4-F3/R3 | Genotyping of *cyp735a4* mutations | ACATGCCGTCCCTCAGCCAC | GGCGTGCTCACCGTATGTCC |
| *proCYP735A3-F/R* | Construction of pCAMBIA1390-proCYP735A3:GUS | CACCACGTCGAACACCTGGTGGCCG | ATGGCCGCCGCCGTCCTCGTC |
| *proCYP735A4-F/R* | Construction of pCAMBIA1390-proCYP735A4:GUS | CACCGCGCACCACCACATGCACACACG | GAGCGACACGAGGACCGCCAT |

**Table S5. List of primers used for quantitative RT-PCR analysis.**

| Gene name | Locus ID | Forward (5' to 3') | Reverse (5' to 3') |
|---|---|---|---|
| *ACT8* | AT1G49240 | AACATTGTGCTCAGTGGTGG | GTGGTGCCACGACCTTAATC |
| *CYP735A3* | LOC_Os08g33300/Os08g0429800 | TACGAGACCGGCAAGAGGAT | GCTCGGAAAATACTGGCTGC |
| *CYP735A4* | LOC_Os09g23820/Os09g0403300 | CGCGCTGATCAAGGAGTTCT | CAGTGTCGTAGCTGGTGTTG |
| *ZIURP1* | LOC_Os03g08010/Os03g0234200 | CACCCTAGGGCTGTCAACTG | GCGAGTGACGCTCTAGTTCT |
| *OsRR1* | LOC_Os04g36070/Os04g0442300 | GTCTCTCGCCTTGTCACCTT | AGAAGCGAAGCACTGATCCT |
| *OsRR2* | LOC_Os02g35180/Os02g0557800 | GTCGCACTACTTCCAGCTCA | GAATTCATGCGCACCACAGG |
| *OsRR4* | LOC_Os01g72330/Os01g0952500 | TGAGAATGTGCCTGCAAGGAT | CAGCGAGCTTGACAGGTTTC |
| *OsRR6* | LOC_Os04g57720/Os04g0673300 | GTGGTGATCATGTCGTCGGA | ATCTGATACGGCTGCAGAGC |
| *OsRR9* | LOC_Os11g04720/Os11g0143300 | CTCTGGAGTTCTTGGGGCTC* | CCAGGCATGCAGTAGTCTGT* |
| *OsRR10* | LOC_Os12g04500/Os12g0139400 | | |
| *OsRR21* | LOC_Os03g12350/Os03g0224200 | GAGTCACAGCATTGGAAGCAG | TGTTTCCAGTGATTACCCTACTTA |
| *OsRR23* | LOC_Os02g55320/Os02g0796500 | CAGACTACAGAGGGATGGCG | AAATGGTGGAGGGCAACAGA |
| *OsSIPP2C1* | LOC_Os09g15670/Os09g0325700 | GTCACCCAGCTGATGCTGTA | CACTGAACTTGGTTAATTCAGGGA |
| *OsGH3.2* | LOC_Os01g55940/Os01g0764800 | GCAAGGGGCTCTACTTCCTG | GTAGTAGCTGGTCAGCACCG |
| *OsIPT4* | LOC_Os03g59570/Os03g0810100 | GTAGATCTCGAGGTGCTCCG | AGATGCCCCTGGAGTAGTCG |
| *OsNIA1* | LOC_Os08g36480/Os08g0468100 | CGTGGACCGTCGATGTGAC | CATGTTCTGCTCCTTGCGG |

*, these primers cannot distinguish *OsRR9* and *OsRR10*
